# Between-subject prediction reveals a shared representational geometry in the rodent hippocampus

**DOI:** 10.1101/2020.01.27.922062

**Authors:** Hung-Tu Chen, Jeremy R. Manning, Matthijs A. A. van der Meer

## Abstract

The rodent hippocampus constructs statistically independent representations across environments (“global remapping”) and assigns individual neuron firing fields to locations within an environment in an apparently random fashion, processes thought to contribute to the role of the hippocampus in episodic memory. This random mapping implies that it should be challenging to predict hippocampal encoding of a given experience in one subject based on the encoding of that same experience in another subject. Contrary to this prediction, we find that by constructing a common representational space across rats in which neural activity is aligned using geometric operations (rotation, reflection, and translation; “hyperalignment”), we can predict data of “right” trials (R) on a T-maze in a target rat based on 1) the “left” trials (L) of the target rat, and 2) the relationship between L and R trials from a different source rat. These cross-subject predictions relied on ensemble activity patterns including both firing rate and field location, and outperformed a number of control mappings, such as those based on permuted data that broke the relationship between L and R activity for individual neurons, and those based solely on within-subject prediction. This work constitutes proof-of-principle for successful cross-subject prediction of ensemble activity patterns in the hippocampus, and provides new insights in understanding how different experiences are structured, enabling further work identifying what aspects of experience encoding are shared vs. unique to an individual.

## Introduction

A fundamental challenge faced by any memory system is how related experiences should be organized – storing the details of each individual experience preserves potentially valuable details, but is storage-inefficient and hampers generalization, whereas treating all experiences as the same risks ignoring potentially important differences^1^. For instance, learning the common spatial features of different floors in the same building makes it possible to predict the layout of a not-yet-visited floor (“similar to the others”); at the same time, each floor also has unique features, such as the location of a specific colleague’s office, that do not generalize. Thus, memory systems need to balance pattern-completion (treating a new observation the same as a previous one) and pattern-separation (keeping similar observations as distinct).

The rodent hippocampus is a model system for studying the neural basis of these processes. Strikingly, the hippocampus can construct statistically independent representations across environments (“global remapping”)^2–5^ and assigns individual neuron firing fields to locations within an environment in an apparently random fashion^6,7^. Similarly, “engram” studies suggest that the population of neurons allocated to a given experience is determined by a competition based on randomly fluctuating excitability levels among eligible neurons^8^. Although there are also examples of hippocampal cells whose firing properties are tied to a particular stimulus feature (e.g. reward^9^) and therefore transfer across different environments, the received wisdom is that those cells that do change their firing fields between environments or across different regions of the same environment, do so randomly^10^.

Remapping studies to date have been limited to within-subject comparisons, but it is possible in principle that what appears random within a single subject in fact obeys a common rule that is shared across subjects. Consider how two related experiences such as running the left (L) and right arms (R) of a T-maze may be encoded in a population of hippocampal neurons (Figure S1). The correlation between L and R activity on a cell-by-cell basis may be zero, but still obey an underlying structure. For instance, cells that tend to fire at the start of L may be more likely to fire at the end of R (Figure S1a), or more realistically, how place fields shift between L and R depends on both their location, firing rate, and relationship to other cells (Figure S1b). If such a rule were to exist, it should be possible to predict, across subjects, what R activity of a target subject looks like, based on (1) that subject’s L activity and (2) the relationship between L and R activity found in a different “source” subject. Although there is no way to predict how two different subjects encode a given experience L (especially when sampling randomly from different numbers of neurons that are not uniquely identifiable across subjects as in e.g. *C. elegans*), the *relationship* between how two different experiences L and R are represented may be conserved across subjects.

Such a *representational geometry* has been demonstrated in a number of brain regions in human cognitive neuroscience studies that use fMRI^11–14^, but cross-subject prediction of this kind has not yet been applied to ensemble recording data in the rodent hippocampus. If (re)mapping in the rodent hippocampus were to show a shared representational geometry, this would not only challenge a long-held dogma about the randomness of place cell allocation, but potentially also open up novel lines of research that can elucidate the algorithmic basis of memory assignment and generalization in a wide variety of settings, while creating a bridge between rodent neural data and human fMRI work.

## Results

The overall goal of this study is to determine if we can predict how hippocampal place cells in a “target” subject encode a particular experience based on two ingredients: (a) knowledge of how the target subject encodes a distinct but related experience, and (b) how a different, “source” subject encodes the same two experiences. To make this prediction possible, we first align the data from source and target subjects in a common representational space, as we describe below.

We operationalize this overall idea using data from T-maze tasks, in which rats run along the left and right arms of the maze to form the two related experiences under study. Specifically, we can describe hippocampal activity on this task as two subject-specific matrices with time as the horizontal dimension, and neuron as the vertical dimension (Figure 1a, leftmost column); one matrix describing the average activity for left trials (L), and another matrix for right trials (R; see Figure S2a for a description of how this input data is obtained). We aim to predict the R matrix in the target subject, based on (a) the target’s L matrix and (b) the source’s L and R matrices. Note that the left and right arms differ in a number of respects other than spatial location, so we use these terms here as descriptive labels, rather than as an interpretation about the nature of their neural encoding (see *Discussion*). We first apply principal component analysis (PCA) to each subject’s data, so that neural activity during L and R trials can be visualized as trajectories in a subject-specific reduced space (Step 1 in Figure 1a).

**Figure 1:**
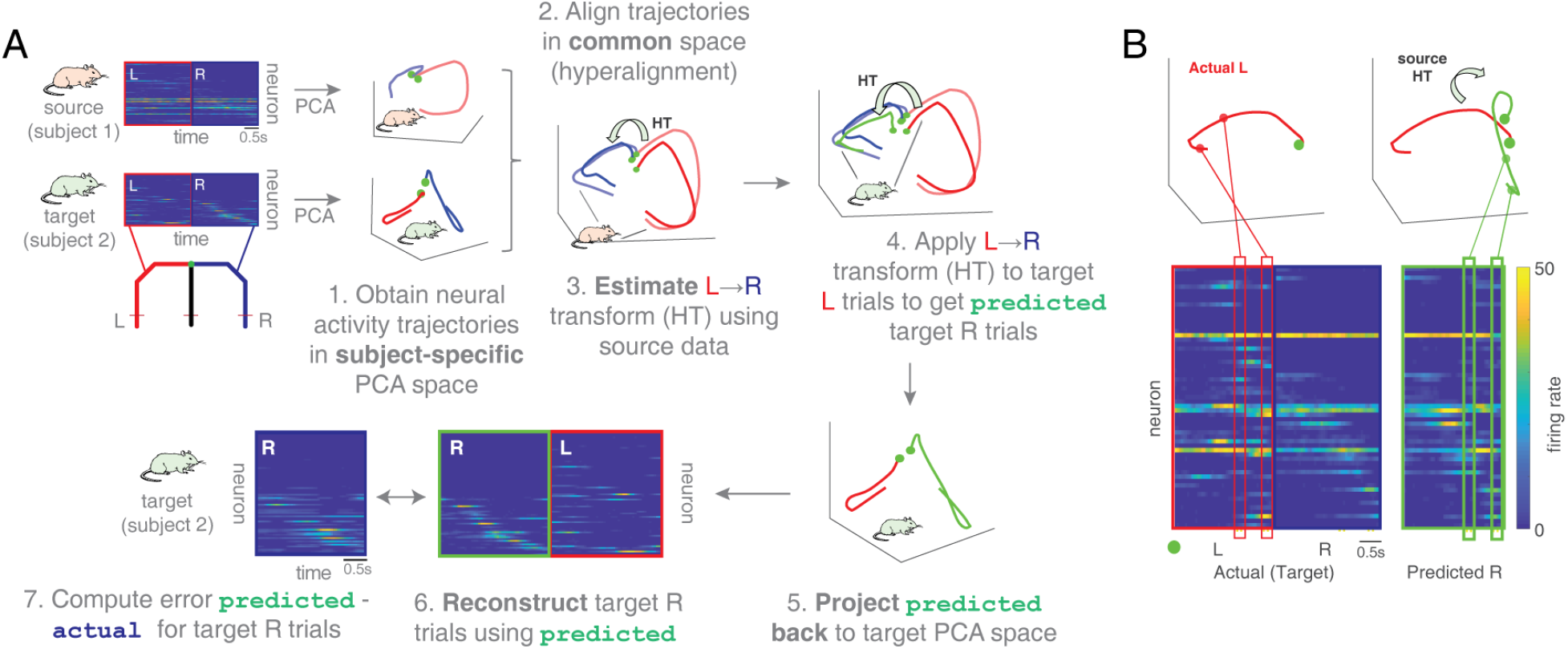
Workflow and example for cross-subject prediction using the hypertransform. **(A)** Our objective is to predict place cell activity on the right arm (R) of a T-maze in a “target” subject, based on (i) place cell activity in the left arm (L) in the target subject, and (ii) L and R place cell activity in a different, “source” subject. These input data are shown in the top left panel: both the source and target subjects have two matrices each that describe, for each recorded neuron, how its activity varies during left and right trials. Neurons are ordered according to their peak firing time on the R arm. Note that although the number of time bins is the same across subjects, the number of recorded cells may be different. Therefore, the first step of the analysis workflow is to apply principal component analysis (PCA), resulting in neural activity trajectories for left and right trials (red and blue, respectively) in each subject’s own PCA space. Three principal components are shown here for display purposes, but in the main analysis 10 PCs were used. Next, these neural activity trajectories are mapped into a common space using a “hyperalignment” procedure that minimizes the Euclidean distance between the L and R trajectories across subjects (step 2, see *Methods* for details). In this common space, a Procrustean transformation^15^ (HT in step 3) is derived that maps L to R trajectories for the “source” subject (step 3), which can then be applied to the L trajectory of the “target” subject (step 4) to obtain its predicted R trajectory in the common space (step 5). This predicted R trajectory is then projected back to the “target” PCA space using the inverse of the matrix used in step 2 (step 6) and expanded back into the target’s original neuron space (step 7). Finally, the predicted R neural activity is compared to the actual R activity to yield an error measure (step 8). The diagonal pattern apparent in the predicted data indicates similarity between actual and predicted R, although differences are also visible. **(B)** Close-up of example target L and R activity matrices (“Actual”, neurons ordered by temporal fields on R) with example predicted R activity derived from a different source session. Each matrix column describes ensemble neural activity for a single time bin, and maps to a corresponding point on a neural activity trajectory in common space (top panels). The hypertransform (HT) is a mapping from L to R activity that operates on these (aligned) ensemble activity vectors. As a result, the R prediction for any given cell is not only based on that cell’s L activity alone, but also depends on the activity of the other cells at that time; see Figure 2 for further examples and a more detailed explanation.

Next, the key step for cross-subject prediction employs a procedure from human cognitive neuroscience, “hyperalignment”^11,16^, that projects each subject’s idiosyncratic neural activity into a common space that minimizes the Euclidean distance between neural activity trajectories (Step 2 in Figure 1a; see also Figure S1a for a schematic). Working in this common space, we can identify the relationship between L and R activity in the source subject, and express this relationship as a transformation matrix (“hypertransform”) which can be applied to the target subject’s L trials to obtain a predicted R trajectory 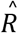. This predicted trajectory is then projected back to the target-specific neural space to obtain a prediction which is compared to the actual data (final step in Figure 1a). Figure 1b shows in more detail an example prediction alongside the actual target activity for a different session; note the overall correspondence between predicted and actual activity (see Figure 2 for more examples).

**Figure 2:**
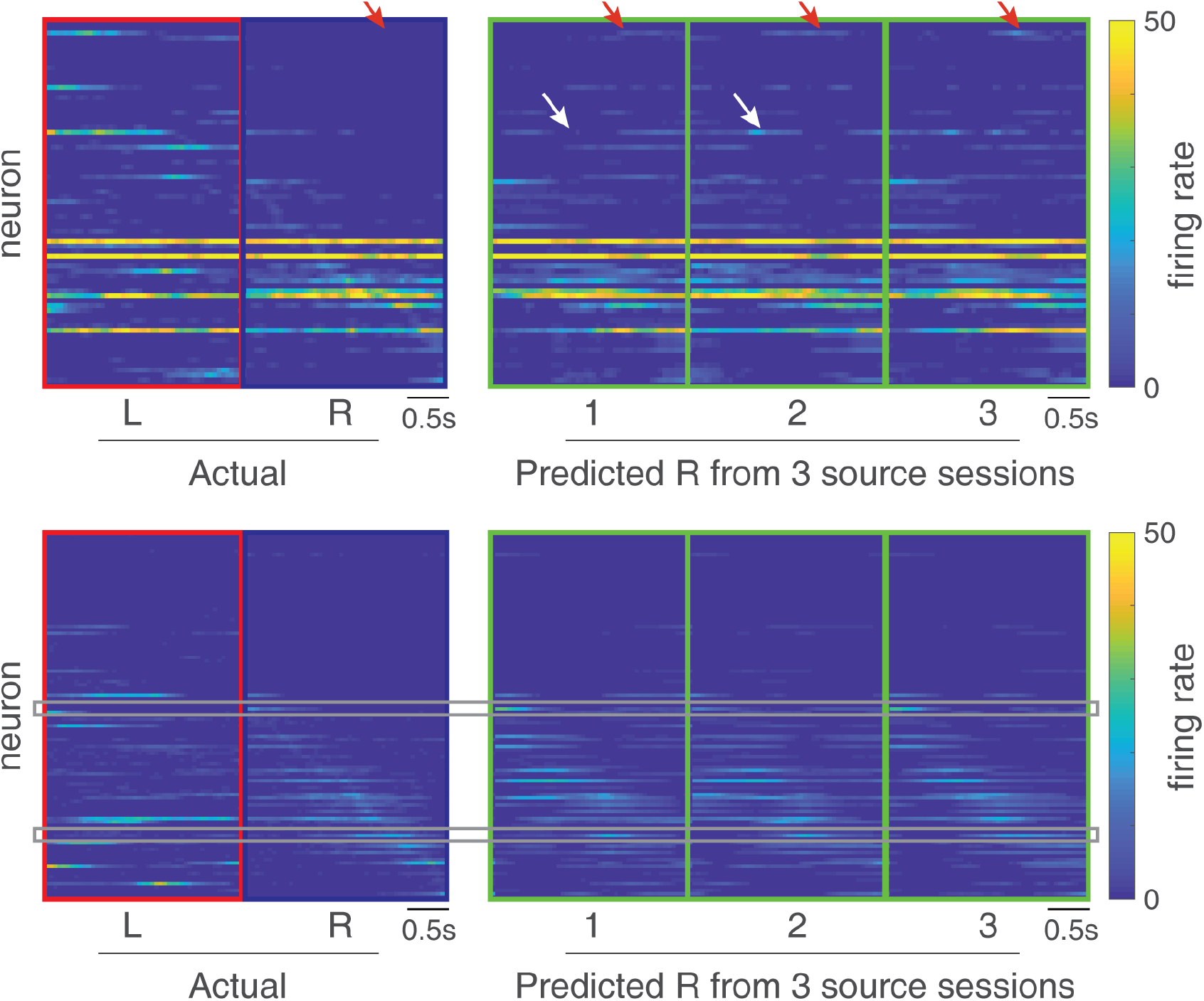
Example target L and R activity matrices with hyperaligned predictions. Two example target L and R activity matrices (“Actual”, neurons ordered by time of maximum activity on R) with hyperaligned predictions obtained from three source sessions (“Predicted”) following the procedure in Figure 1. In general, for each session, the predicted and actual activity are clearly related, as is apparent from the overall diagonal pattern in the Predicted matrices indicating agreement with the Actual R data (we quantify this in insets of Figures 3 and 4 and associated analyses). However, clear deviations from the actual data are also visible, for instance in predicting fields that do not exist in the actual data (red arrows; compare actual and predicted R), and predicting fields in incorrect locations (appearing away from the diagonal, e.g. white arrow for source session 2). Comparing between different source sessions (1, 2, 3) the predictions derived from all source sessions share an overall similarity, but there are also differences, typically in the specific locations of predicted place fields in a subset of neurons (compare the two white arrows, for instance). A further point to note is that even for some cells that do not have a field on L, we can correctly predict where this cell will have a field on R (gray rectangles in lower panel). In general, this occurs because the hyperalignment step rotates, reflects and translates the source and target data to minimize the distance between them, such that neurons that change similarly across subjects become aligned. In more detail, we can ask, how is it that even though the L activity is the same at different time points, the predicted R is different over time? This occurs because the mapping from L to R at each time point depends on the activity of all the other cells in L at that time point. Next, how is it that for a given time point, even though the L activity is the same for two different cells, the predicted R is different for those cells? This occurs because different cells have different loadings on the principal components that are hyperaligned. In other words, cross-subject prediction does not apply a fixed rule to the activity of a single cell, but rather applies a mapping that depends on the activity of the other cells in the ensemble.

If a given subject encodes L and R trials independently, then it should not be possible to use one subject’s neural activity for L and R to predict anything about how another subject encodes R trials based on its L trials. On the other hand, if there is some shared structure between subjects in how L and R trials are encoded, then cross-subject prediction should perform better than chance. To test this idea, we compare the prediction of a target subject’s R trials to shuffled controls. In the first, “row shuffle”, we obtain a distribution of chance predictions for each source-target pair, based on breaking the relationship between L and R trials in the source subject by randomly permuting the rows of the R matrix (see Figure S2b for a schematic). Based on this chance distribution, we define three metrics: (1) a z-score of the actually observed error compared to chance, (2) the difference between the actually observed prediction error per cell and the mean of the chance prediction error, and (3) the proportion of chance prediction errors that were lower than the observed error.

We used two different data sets: the first, “Carey” data set^17,18^ is from a T-maze where L and R arms were deliberately furnished with distinct surface colors and textures. In contrast, the second, “Gupta” data set^19,20^ used a T-maze whose arms had similar surfaces. Starting with the Carey data, we found that the hypertransform (HT) prediction of R trials in the target subject was better than chance overall for all metrics used (Figure 3, top and middle rows; green “HT” bars; z-score: p < 0.001 for Wilcoxon signed rank test vs. 0; raw error: p < 0.001). Cross-subject prediction of R activity was better than chance even when the R data was withheld entirely from the hyperalignment step (see *Methods* for details on the withholding procedure; Figure 4a, top row; p < 0.001 for HT vs. 0), when L and R activity was expressed as tuning curves in space rather than in time (Figure 4c, middle row; p < 0.001 for HT vs. 0), and when putative interneurons were removed (Figure S3a). We found similar results for the Gupta data, with the HT prediction consistently better than chance (Figure 3, bottom row; p < 0.001 for HT vs. 0). These results demonstrate that the relationship between L and R trials is not random across subjects.

**Figure 3:**
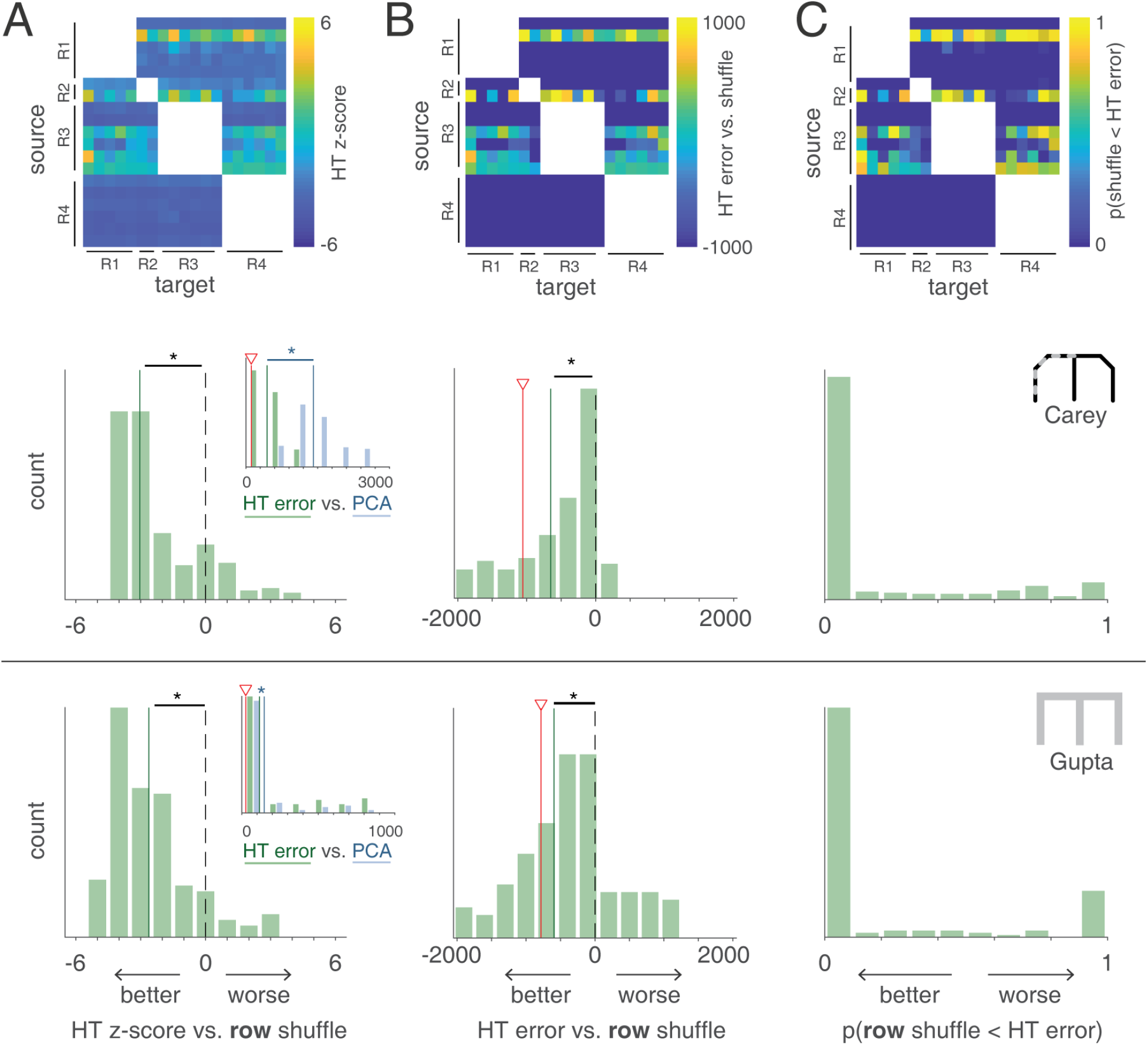
Cross-subject prediction of R trials of a “target” subject based on how a “source” subject encodes L and R trials outperforms prediction based on shuffled source data. For each source-target pair in the Carey data, we computed a z-score of the actual observed error between predicted and actual R trials (based on the hypertransform procedure, “HT”) compared to a shuffled distribution in which the R rows of the source subject were randomly permuted. Thus, lower z-scores indicate lower error and therefore better prediction than chance. Across all source-target pairs, this z-scored error varied depending on the pair used (column **A**, top row), but was lower than chance overall, as indicated by a shift in the z-score histogram relative to 0 (“HT” green bars in column **A**, middle row; median: −3.10 +/− 0.54, SEM across unique source-target pairs, p < 0.001 for Wilcoxon signed rank test vs. 0). Cross-subject predictions based on the L-R transform in common space (“hypertransform”) outperformed predictions based on the L-R transform in PCA space (“PCA” blue bars in column **A**, middle row, inset; see *Methods* for details; HT < PCA: 99.62% of source-target pairs, p < 0.001 for binomial test) in terms of raw error between predicted and actual R trials (median of HT raw error: 443.73 +/−62.44, PCA: 1409.14 +/−204.22,split-half: 130.99 +/−46.72, red triangle in inset; mean squared error per cell, see *Methods* for details). Next, we applied the same analysis to a different data set (“Gupta”, bottom row, in which the L and R maze arms were more similar to each other than in the “Carey” data), and found that the hypertransform prediction was again significantly better than chance (“HT”; median: −2.59 +/− 1.44, p < 0.001, bottom row) and better than PCA-only (HT < PCA: 79.58% of source-target pairs, p < 0.001; column **A**, bottom row, inset; median of HT raw error: 115.26 +/− 142.05, PCA: 121.42 +/− 208.23, split-half: 37.39 +/− 10.01). Columns **B** and **C** use the same layout as column **A**, but using different metrics to describe prediction accuracy. **B** uses the raw error (between predicted and actual R neural activity; lower error/negative indicates better prediction) compared to the mean of the shuffle distribution, and **C** uses the proportion of the shuffle distribution with smaller error than the actually observed error (lower proportions indicate better prediction). For the raw error measure, the HT prediction was better than chance in both the Carey data (column **B** middle row; median: −652.87 +/− 486.05, p < 0.001 for HT vs. 0; red line indicates split-half prediction) and the Gupta data (**B** bottom row; median: −594.62 +/− 735.37, p < 0.001 for HT vs. 0).

**Figure 4:**
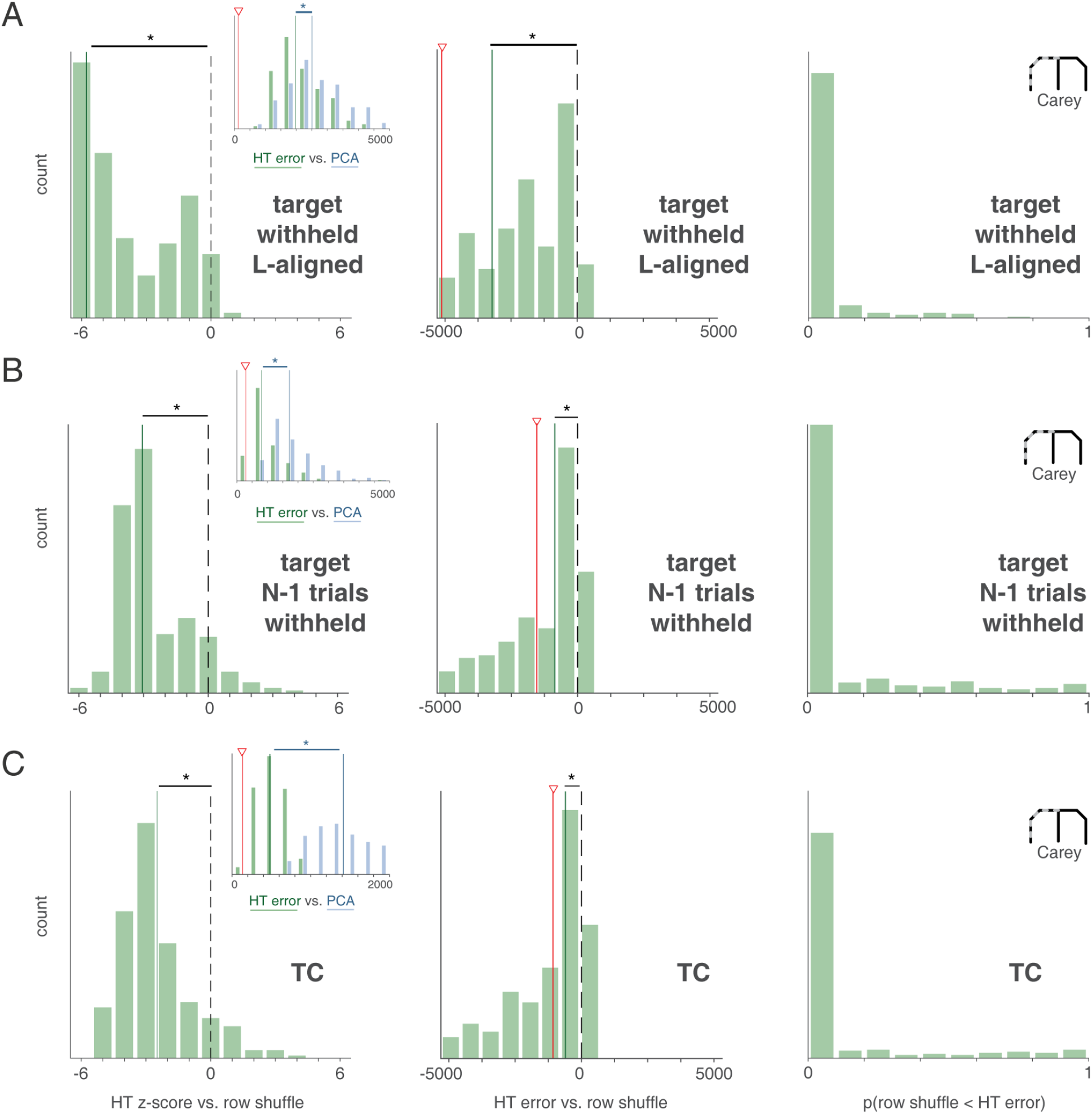
Better-than-chance cross-subject predictions can be observed even when the to-be-predicted target data was withheld, and spatial turning curves were used. **A**: Histogram of three cross-subject prediction metrics: z-scores of actual observed error compared to the distribution of shuffle predictions (left column), raw prediction error compared to the mean of the shuffle distribution (middle), proportion of the shuffle distribution whose error was smaller than actual observed error (right). For all metrics, lower numbers indicate better cross-subject predictions. Even when the R activity of the target subject, which is the activity to be predicted, is withheld from the hyperalignment procedure, the hypertransform (HT) prediction is significantly better than chance (median: −6.03 +/− 0.84, SEM across unique source-target pairs, p < 0.001 for Wilcoxon signed rank test vs. 0; left column) and better than PCA-only (HT < PCA: 70.77% of source-target pairs, p < 0.001 for binomial test; left column, inset; median of HT raw error: 1962.93 +/− 209.86, PCA: 2505.85 +/− 287.00, split-half: 114.05 +/− 17.48, red triangle in inset). **B:** Histogram of three cross-subject prediction metrics as in **A**. Different from **A**, one randomly-chosen trial of target R was used for alignment, with the average of rest R trials withheld as the activity to be predicted. The HT prediction is again better than chance (median: −3.03 +/− 0.51, p < 0.001 for HT vs. 0; left column) and significantly better than PCA-only (HT < PCA: 91.92% of source-target pairs, p < 0.001; left column, inset). Note that the HT prediction was substantially improved by including one trial of target R data (median of HT raw error: 809.73 +/− 374.22, PCA: 1698.19 +/− 345.25, split-half: 108.83 +/− 110.81). **C**: Histogram of three cross-subject prediction metrics as in **A** and **B** for neural activity matrices calculated as a function of locations (turning curves; TC) instead of time (see Figure S2 and *Methods* for details) were used. The cross-subject predictions are significantly better compared to shuffles (median: −2.97 +/− 0.46, p < 0.001 for HT vs. 0; left column) and significantly better than PCA-only (HT < PCA: 100% of source-target pairs, p < 0.001; left column, inset; median of HT raw error: 499.05 +/− 50.22, PCA: 1469.76 +/− 170.06, split-half: 83.86 +/− 64.48), suggesting time and location yield similar results (compare with Figure 3).

To test if the hyperalignment step of the prediction procedure is important, rather than some other part of the workflow, we repeated the analysis with the hyperalignment step left out (i.e. we applied the L-R transform obtained from the source subject’s PCA space to the target subject’s PCA space, “PCA-only”; see Methods for details). The HT-based prediction outperformed the PCA-only prediction both for the Carey data (blue “PCA” bars in Figure 3a inset, middle row; p < 0.001, binomial test) and the Gupta data (bottom row; p < 0.001, binomial test), although for the Gupta data the difference was noticeably smaller (we investigate this further below). Again, the HT predictions outperformed PCA-only predictions when spatial tuning curves were used (Figure 4c) and with putative interneurons removed (Figure S3a); to facilitate visualization of the results in the analyses that follow, we used the interneuron-removed data. Although the HT prediction still outperformed PCA-only when target-R data was withheld entirely, the HT prediction was much better (i.e. closer to split-half prediction) if at least one trial of target R was included for alignment (Figure 4a and b; insets). Crucially however, R data cannot be predicted better than chance even when it is included in the alignment step when in fact no underlying L-R relationship exists, as we verify below with simulated data.

A possible trivial explanation for the better-than-chance cross-subject prediction is that rats represent experiences in L and R similarly, so that a simple duplicate of L activity forms a reasonable prediction of R activity. To test if L-R correlations at the cell-by-cell level (i.e. row-wise correlations, Figure S2c) underlie the cross-subject prediction results in Figure 3, we compared cross-subject predictions based on the hypertransform (L-R mapping in common space) with those based on the *identity transform*: a within-subject prediction that simply takes a duplicate of the L trajectory in common space and uses it as the prediction for R. For the Carey data set, the cross-subject HT prediction was significantly better than that based on the identity transform (ID; left panel in Figure 5a; p < 0.001, binomial test), demonstrating that the better-than-chance prediction of R is not due to cell-by-cell correlations with L activity. In contrast, for the Gupta data, the HT prediction did not outperform the identity prediction (right panel in Figure 5a), suggesting that the better-than-chance cross-subject prediction for this data set (Figure 3, bottom row) can be attributed to row-wise correlations between L and R activity.

**Figure 5:**
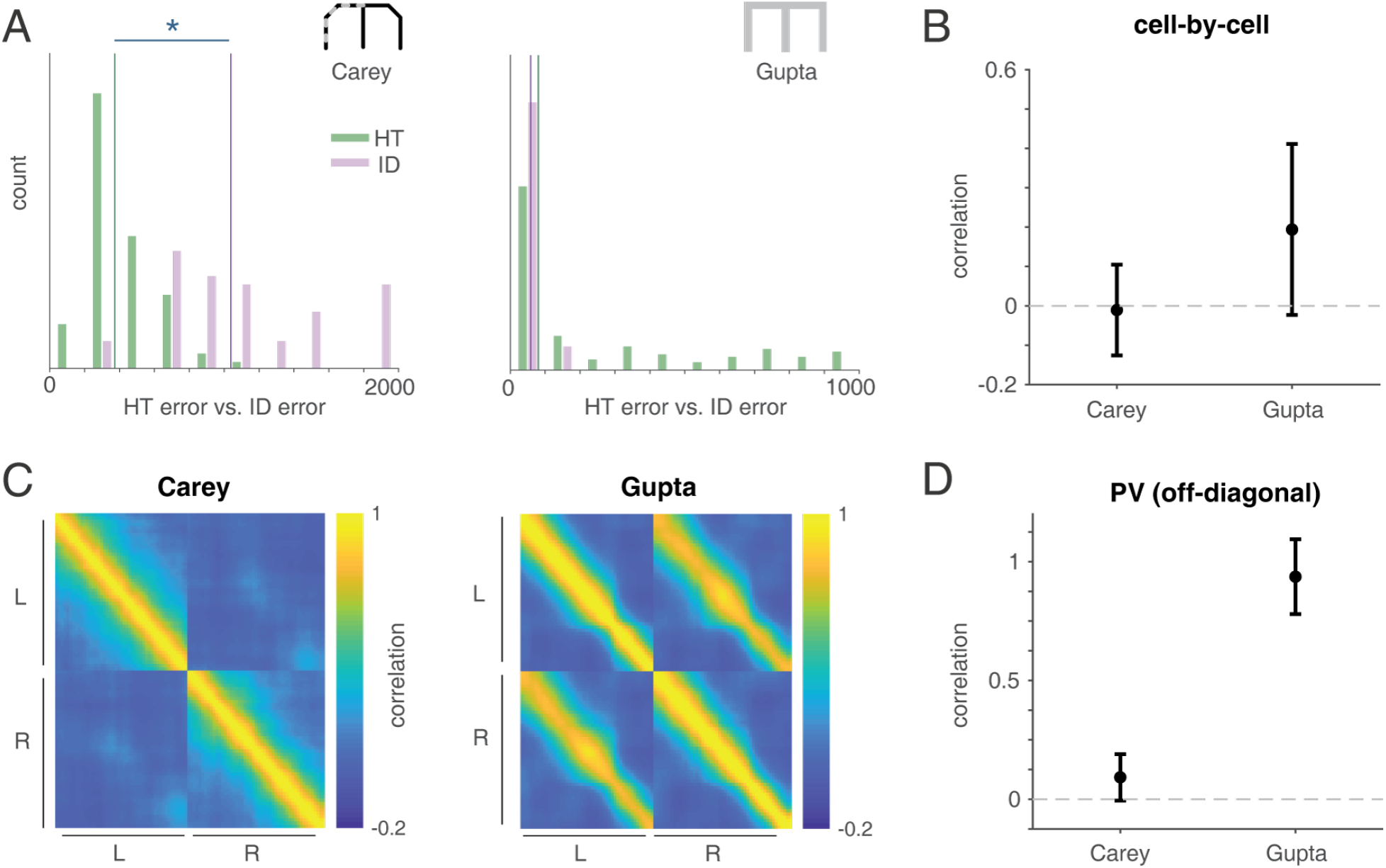
Cross-subject prediction outperforms within-subject prediction only in the absence of cell-by-cell correlations. **A**: Comparison of cross-subject prediction error (“hypertransform”, green bars; HT) with within-subject prediction error (“identity transform”; blue bars, ID) for two different data sets. In the “Carey” data (left panel) the left and right arms of the maze had different texture and color patterns; in the “Gupta” data (right panel) the two maze arms were identical. For the Carey data, cross-subject prediction was significantly better than within-subject prediction (HT < ID: 100% of unique source-target pairs, p < 0.001 for binomial test) whereas for the Gupta data, this difference was not significant. **B**: Cell-by-cell correlations of firing rates between L and R arms (i.e. row-wise correlations of the L and R matrices in Figure S2c), were significantly higher in the Gupta data compared to the Carey data (Gupta: median r = 0.21 +/− 0.27, SEM across subjects, Carey: median r = −0.02 +/− 0.12, Wilcoxon rank sum test, p < 0.001). Cell-by-cell correlations in the Carey data were not significantly different from 0 (p = 0.44 for Wilcoxon signed rank test vs. 0). **C**: Population vector (PV) correlations between ensemble activity at each time point and every other time point, i.e. column-wise correlations of the L and R activity matrices, averaged across sessions. Both Carey and Gupta data sets show high correlations around the diagonal, indicating an overall autocorrelation in time; however, the Gupta data additionally shows high off-diagonal correlations between L and R which are barely visible in the Carey data. **D**: Quantification of the median PV correlation between L and R (i.e. the values along the diagonal of the lower left quadrant in **C**). For Gupta data, this correlation is remarkably high (median r = 0.95 +/− 0.16) whereas for Carey data, it is significantly lower (median r = 0.09 +/− 0.10, p < 0.001 for Wilcoxon rank sum test) but significantly different from 0 (p < 0.001 for Wilcoxon signed rank test vs. 0), consistent with previous reports of global remapping^2,4^. The above results explain why the HT is not needed for Gupta data to achieve better than chance predictions (as shown in Figure 3, bottom row): L and R activity is sufficiently similar such that the L trajectory alone in either PCA or common space can predict R activity.

To test this idea, we investigated the correlation structure between the L and R firing rate matrices using two different measures: the cell-by-cell (row-wise) correlation, averaged across all cells, and the column-wise population vector (PV) correlation (averaged across sessions, see Figure S2 for schematic; note that in order to compute these measures, putative interneurons were removed from the data; see Figure S3a and Methods). The cell-by-cell firing rate correlations between L and R arms were significantly more correlated in the Gupta data compared to Carey data (Gupta: median r = 0.21 +/− 0.27, Carey: median r = −0.02 +/− 0.12, SEM across subjects; p < 0.001, Wilcoxon rank sum test; Figure 5b). Similarly, PV correlations in the Gupta data showed high off-diagonal values between L and R, which were barely visible in the Carey data (Figure 5c-d). The low correlation values observed in the Carey data are consistent with those previously reported and characterized as global remapping^2,4^, whereas the Gupta correlations are strikingly high, indicating the presence of “symmetric” cells with similar firing patterns on the L and R arms. High L-R correlations in the Gupta data imply that the cross-subject (hypertransform) method cannot outperform the already very good prediction based on within-subject correlations, whereas for the nearly uncorrelated Carey data, there is room for cross-subject prediction to improve.

Importantly, the comparison between the two data sets suggests that the cross-subject prediction on Carey data is not the result of within-subject correlations -- because, if it were, then the HT prediction would be similar to the ID prediction. So, if cross-subject prediction for the Carey data is not simply a consequence of within-subject correlations between L and R, what *is* the prediction based on? In other words, can we identify what features of the L-R relationship are generalizable across subjects without appearing as within-subject correlations between L and R activity? To address this question, we generated synthetic neural activity matrices using 1-D Gaussians with three parameters: time, peak firing rate (FR) and width. Specifically, three simulated data sets captured different potential place cell properties: (1) each neuron has an independent probability of having a firing field on L and R, and all parameters are randomly and independently chosen for L and R (**ind-ind**, top row in Figure 6a), (2) if a neuron has a firing field on L, it does not have a field on R (and vice versa), and the parameters of the field are chosen randomly (**x-or**, second row) and (3) each neuron has an independent probability of having a field on L and R, but if a cell has a field in both, all three parameters are the same (**ind-same-all**, third row). For all these scenarios, we generated synthetic data matching the number of recording sessions and the number of neurons recorded in the Carey data, and applied exactly the same analysis procedure.

**Figure 6:**
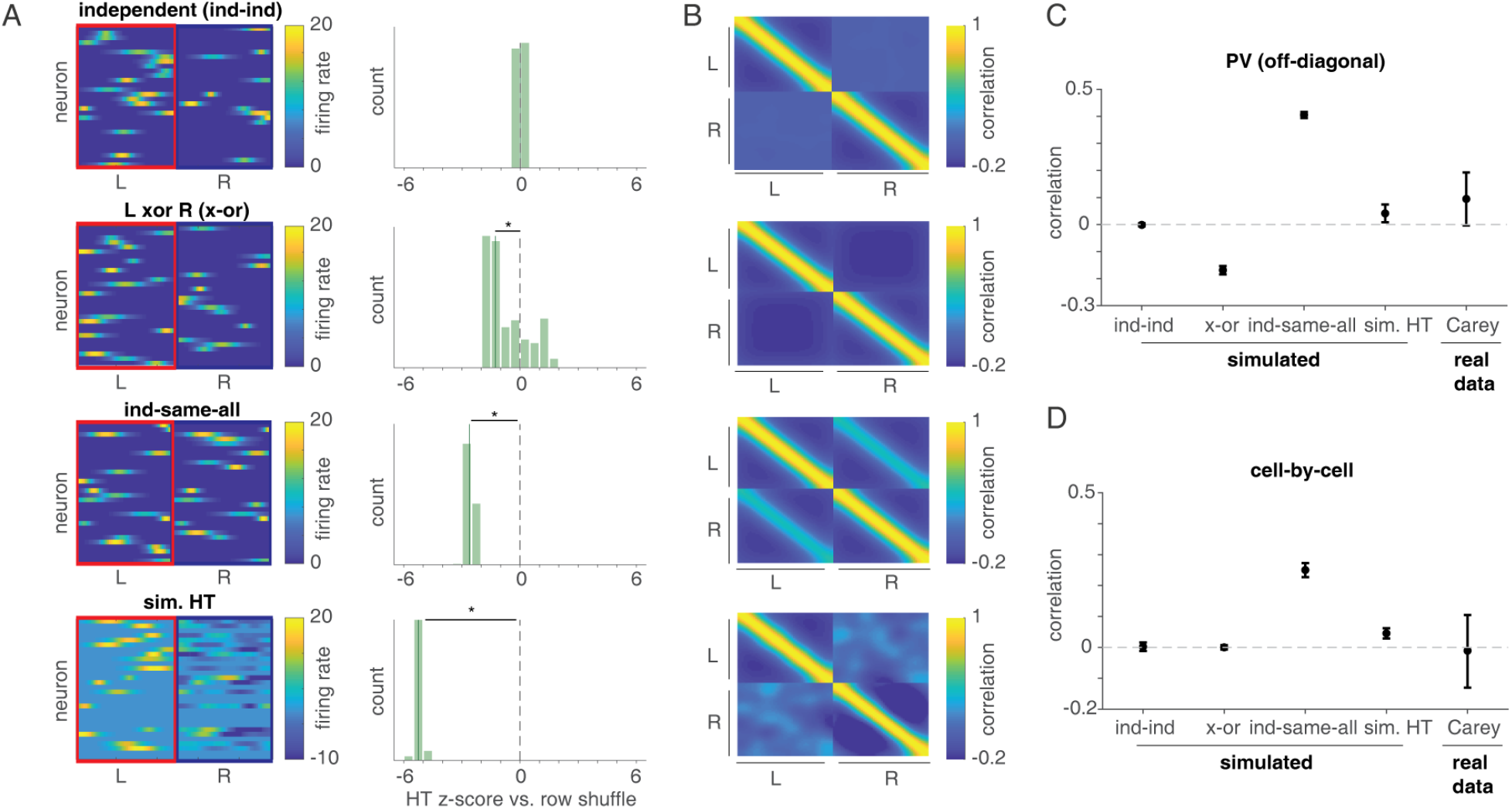
Simulations demonstrate that representational geometry, but not simple rules such as exclusive-or and firing rate correlations, result in cross-subject prediction while being consistent with the data. **A**: Example L and R activity matrices (left column) and histogram of z-scores of cross-subject prediction compared to the distribution of shuffle predictions (right column; z-scores lower than zero indicate better-than-chance predictions) of four simulated data sets: (1) neurons have a fixed, independent probability (0.5) of having a 1-D Gaussian place field on L and/or R, with the three parameters of time, peak firing rate (FR) and width randomly and independently chosen for L and R (**ind-ind**, top row), (2) neurons only have a field on *either* L or R but not both, and parameters of the field are chosen randomly as in (1) (**x-or**, second row), (3) neurons have a fixed independent probability of having L and R fields as in (1) but with the additional constraint that neurons with both L and R fields must have the same three parameters (**ind-same-all**, third row) and (4) the activity on L is simulated by assigning each neuron an independent probability of having a field whose parameters are randomly chosen, then the activity on R is obtained by applying a L-R transform (hypertransform, HT) from the Carey data to the simulated L activity (**sim. HT**, last row). As expected, in the **ind-ind** (independent) case, cross-subject prediction is not possible (z-score vs. chance, median: −0.01 +/− 0.03, SEM across unique source-target pairs, p = 0.70 for Wilcoxon signed rank test vs. 0). In contrast, **x-or**, **ind-same-all** and **sim. HT** all show better-than-chance cross-subject predictions (median: −1.26 +/− 0.30 for **x-or**, median: −2.62 +/− 0.06 for **ind-same-all**, median: −5.23 +/− 0.04 for **sim. HT**, all p < 0.001 for Wilcoxon signed rank test vs. 0), indicating that if there is a non-random L-R relationship in the underlying data, the hypertransform procedure can exploit it. Note that although this synthetic example shows substantially negative predicted firing rates, negative firing rates, when they did occur, tended to be much smaller for predictions using actual data. **B**: Population vector (PV; column-wise) correlations between ensemble activity at each time point and every other time point of the L and R activity matrices. Only **ind-same-all** shows high off-diagonal correlations between L and R, resembling the Gupta data set in which L and R arms were identical and firing activity on both arms is highly correlated (compare with Figure 5c). In **x-or**, off-diagonal correlations are slightly negative. **C**: Quantification of the median PV correlation between L and R (i.e. the values along the diagonal of the lower left quadrant in **B**). PV correlations between L and R in **ind-ind** are zero, but L and R are positively correlated in **ind-same-all**(median r = 0.40 +/−) since for every time point in L where there is a field, the same ensemble activity appears at the same time point in R with probability 0.5. **X-or** shows a negative correlation (median r = −0.18 +/− 0.02) because for every time point in L where there is a field, the same ensemble activity would deterministically be absent in R, and vice versa. None of these simple rules are consistent with the PV correlation found in the Carey data; however, **sim. HT** does show similar correlations as the data (median r = 0.04 +/− 0.03 for **sim. HT** and median r = 0.09 +/− 0.10 for Carey). **D**: Cell-by-cell (row-wise) correlations of L and R show that **ind-same-all** is more correlated than the data, whereas **sim. HT** yields similar correlations (median r = 0.04 +/− 0.02 for **sim. HT** and median r = −0.02 +/− 0.12 for Carey). Thus, taken across panels C and D, simple rules (**x-or** and **ind-same-all**) are inconsistent with the data, but the representational geometry embodied in the **sim. HT** yields correlations that are similar to the data.

The independent (**ind-ind**) simulation serves as a crucial sanity check to verify that our cross-subject prediction procedure cannot exploit shared structure where none exists; as expected, cross-subject prediction was not different from chance in this scenario (Figure 6a, right column; p = 0.92 for Wilcoxon signed rank test vs. 0). In contrast, both **x-or** and **ind-same-all** showed better-than-chance cross-subject prediction (both p < 0.001). If the **x-or** or **ind-same-all** rules are potential explanations for better-than-chance predictions in Carey data, we should see zero cell-by-cell correlations and low PV correlations between L and R in these two simulations as observed in the Carey data (Figure 5b-d). However, the **x-or** scenario shows negatively correlated PV correlations inconsistent with the Carey data (Figure 6b-c; r = −0.17).

The **ind-same-all** rule shows both high cell-by-cell correlations (Figure 6d; r = 0.25) and high PV correlations (r = 0.41), which is again inconsistent with the Carey data, but more in line with the Gupta data (compare Figure 5b-d). In further simulations, we separately investigated the role of each parameter of the 1-D gaussian place fields (time, FR and width) as a potential explanation for cross-subject prediction; only when the same time is shared across L and R are the cross-subject predictions better than chance (**ind-same-time**, top row in Figure S4; p < 0.001). However, as in the main analysis (Figure 5b) the cell-by-cell correlations are inconsistent with those observed in the Carey data. Thus, although both rules like x-or and same-time can support cross-subject prediction, the cell-by-cell correlations in the Carey data are different and therefore these simple rules cannot be the full explanation.

If these predetermined rules cannot be the source of cross-subject prediction in the Carey data, what can? We first sought to rule out a number of possible low-level explanations, verifying that differences between source and target sessions in running speed (Figure S5a) and average firing rates (Figure S5b) do not explain cross-subject prediction performance. Plotting cross-subject prediction error as a function of time did not highlight any particular areas as being especially well or poorly predicted (Figure S6c) ruling out explanations based on shared cues such as the choice point or the existence of a reward site at the end of each arm. One more subtle possibility is a *single-cell mapping rule* such as a cell with a field at the start of L being mapped to the end of R; however, no obvious field mapping pattern between L and R was observed for both actual and predicted data (Figure S7), consistent with the low population vector correlations in the data (Figure 5c-d).

Is there another possibility? The hyperalignment procedure we used to derive the L-R transform (“hypertransform”) was originally developed to capture so-called *representational geometry*: a shared rule that specifies how differences in the *ensemble* neural response to a set of stimuli may be preserved across subjects even though each subject may encode a given stimulus quite differently^11^. To test if such a geometry is consistent with the data, we created another synthetic data set in which the activity on L is simulated by assigning each neuron an independent probability of having a 1-D gaussian place field whose parameters are randomly chosen, and the activity on R is obtained by applying the L-R transform (hypertransform, HT) from the Carey to the simulated L activity (**sim. HT**, bottom row in Figure 6a, see *Methods*). Not only did **sim**. **HT** show significant better-than-chance cross-subject predictions on the simulated data (p < 0.001) but the cell-by-cell (r = 0.04) and PV correlations (r = 0.04) were similar to the Carey data. Thus, unlike the simple x-or and same-parameter simulated scenarios, and unlike single-cell mapping rules, a shared representational geometry is consistent with the correlations observed in the data.

The hypertransform capitalizes on a geometric relationship between ensemble activity patterns, which can be difficult to relate back to the properties of any individual cell. However, some insight can be obtained by separating (1) *mean firing rate* relationships between L and R regardless of their specific firing locations (e.g. “cells with high rates on L tend to have high rates in R”, as suggested by the data in Lee et al. 2019^21^), and (2) *specific firing locations* regardless of mean firing rate differences (e.g. “a cell that was co-active with cells ABC on L will be co-active with cells DEF on R”). To determine the contributions of *mean firing rate* relationships, we applied different firing rate normalization methods to each cell’s L and R activity independently (note that normalizing each cell’s L and R joint activity is not appropriate because it introduces an artificial anticorrelation between L and R that the hypertransform will exploit). L2 normalization preserved a significant but smaller amount of cross-subject prediction compared to the unnormalized data (Figure S3b), while Z-scoring firing rates abolished cross-subject prediction (i.e. did not outperform chance level; Figure S3c, see figure legend for our interpretation of this difference). Reduced cross-subject prediction after firing rate normalization shows that firing rate relationships between L and R are an important component of the prediction. To determine the contributions of specific *firing locations*, we performed an additional shuffle that circularly shifted the R activity by a different random amount for each cell (see Figure S2 and *Methods*). The full (non-shuffled) cross-subject prediction outperformed this circular shuffle both for the Carey data (Figure 7, middle row; p < 0.001 for Wilcoxon signed rank test vs. 0) and the Gupta data (bottom row; p < 0.001 for HT vs. 0). Taken together, these analyses demonstrate that both single-cell firing rate and firing location relationships contribute to cross-subject prediction.

**Figure 7:**
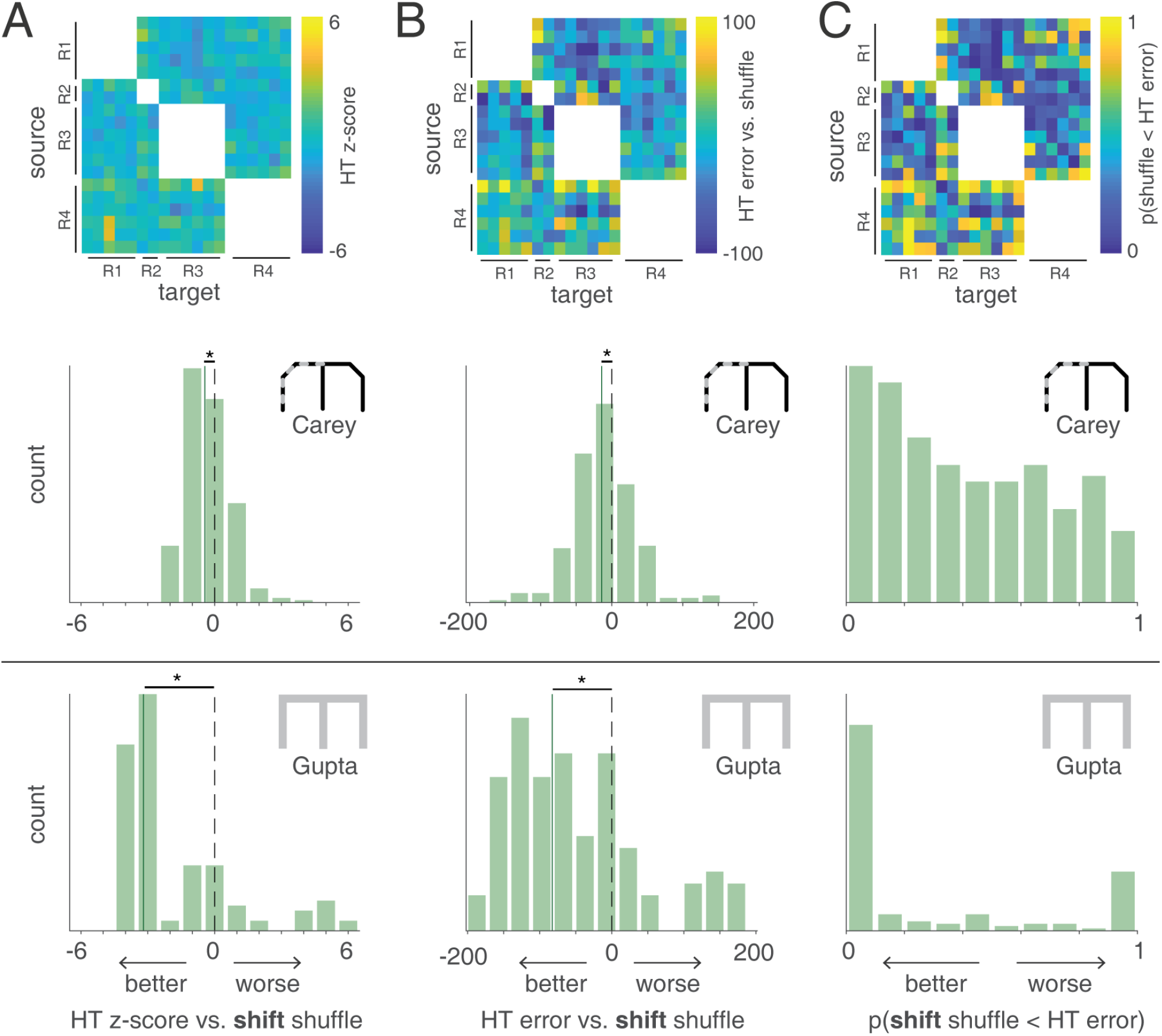
Cross-subject hypertransform (HT) outperforms temporally shuffled source data. For each source-target pair in the Carey data, we computed a z-score of the actual observed error between predicted and actual R trials (based on HT) compared to a shuffled distribution in which each R row of the source subject was *circularly shifted independently by a randomly chosen length*. Lower z-scores indicate lower error and therefore better prediction than chance. Across all source-target pairs, this z-scored error varied depending on the pair used (column **A**, top row), but was lower than chance overall, as indicated by a shift in the z-score histogram relative to 0 (“HT” green bars in column **A**, middle row; median: −0.44 +/− 0.29, SEM across unique source-target pairs, p < 0.001 for Wilcoxon signed rank test vs. 0). Next, the same analysis was applied to a different data set (“Gupta”, bottom row, in which the L and R maze arms were more similar to each other than in the “Carey” data), and found that the HT prediction was again significantly better than chance (“HT”; median: −3.20 +/− 1.13, p < 0.001, bottom row). Column **B** and **C** use the same layout as column **A**, but using different metrics to describe prediction accuracy. **B** uses the raw error (between predicted and actual R neural activity; lower error/negative indicate better prediction) compared to the mean of the shuffle distribution, and **C** uses the proportion of the shuffle distribution with smaller error than the actually observed error (lower proportions indicate better prediction). For the raw error measure, the HT prediction was better than chance in both the Carey data (column **B** middle row; median: −14.12 +/− 12.27, p < 0.001 for HT vs. 0 and the Gupta data (**B** bottom row; median: −206.13 +/− 119.78, p < 0.001 for HT vs. 0).

## Discussion

We have shown that it is possible to predict across subjects, better than chance, how a given experience will be encoded in the hippocampus. In particular, we describe how a “target” subject represents the right arm of a T-maze (R), given (1) how that same subject represents the left arm (L) of the same maze, (2) the relationship between L and R activity in a different “source” subject, by aligning neural activity from both subjects in a common space. In a true predictive version of the analysis, where the R activity of the target subject is withheld from the alignment step, we can still predict R activity better than chance, suggesting this activity is not merely random but has a geometric relationship to L activity that generalizes across subjects. Control analyses based on within-subject prediction and a comparison of the properties of various simulated data sets and shuffles with the real data suggests that this cross-subject prediction is unlikely to be the result of trivial relationships such as cells with symmetric firing fields, fixed field mapping rules like the start of the L being mapped to the end of R, or simple rules such as exclusive-or. Thus, our results imply that the hippocampal encoding of different locations in space, commonly reported to be random *within* subjects^2–4,6,7^, in fact has a shared structure *between* subjects.

There have long been indications in the literature that the space of possible neural activity patterns in the hippocampus is constrained in a number of ways, perhaps most obviously by a continuous attractor-type structure that enforces a single, coherent “hill of activity” to exist within a map (chart) at a time^22^. A different example is how the encoding of a novel experience is constrained by internally generated activity preceding that experience (“preplay”)^23,24^, and most recently, the demonstration that the formation of new place fields in response to optogenetic stimulation is predictable from activity prior to the stimulation^25^. Our findings are congruent with these overall notions, but add a new dimension in its ability to test and reveal structure shared *between* subjects. This approach is attractive because it provides rigorous, quantifiable tests of how generalizable a given model of neural activity is, and because it can provide insights into what is shared and what is unique between subjects. Related work using calcium imaging of CA1 neurons has successfully decoded location and other task variables in one animal even when the decoder is trained on data from another animal^26,27^. Our approach is similar in that it also uses cross-subject prediction, but addresses a different question in that we seek to predict not the decoded location of the animal in task space given a common experience, but the actual neural activity pattern corresponding to a related but different experience.

Unlike in the rodent literature, a substantial number of human fMRI and ECoG studies have used cross-subject prediction^11–13,28,29^. Particularly effective are procedures that do not only align across subjects anatomically (e.g. by mapping each subject to a reference brain) but functionally, i.e. by finding structure in how related experiences are represented, even though across subjects the same experience may be represented very differently. Haxby et al. (2011)^11^ refer to such shared structure as “representational geometry”, an idea implemented here as a rotation and translation (Procrustes transformation) in ensemble neural activity. Because this notion is abstracted away from the raw data and describes a relatively general class of relationships, the question arises what specific features of the data make cross-subject prediction possible. In particular, can this abstract geometric concept be reduced to a simpler set of rules? Hypertransform predictions do not appear to be reducible to single-cell rules, but rather are based on a mapping in ensemble activity space (i.e. how a given L cell’s activity changes between L and R depends on what cells are co-active with it). This idea is illustrated by the examples in Figure 2, and supported more formally by (1) comparison of correlations in the data with simulations (Figure 6), and (2) outperforming both row (firing rate) and shift (firing location) shuffles. We note that the contribution of firing location differences is smaller compared to firing rate differences, as shown by comparing the middle rows of Figure 3 and Figure 7, and by the inability to outperform row shuffles when data is z-scored (Figure S3c; this tends to introduce noise that masks underlying regularities); nevertheless, these analyses demonstrate a contribution of both to an underlying ensemble-level regularity.

What circuit-level or hippocampal input properties are involved in this shared geometry? One possibility is suggested by the relationship between grid cells and place cells: Dordek et al. (2016)^30^ showed that applying nonnegative principal component analysis (PCA) to ensemble activity of place cells yields grid cell-like activity patterns. In addition, grid cell firing patterns were known to remain intact but realign linearly (and differently for different subjects) during global remapping^31^. This suggests that our hypertransform procedure may rely on a common coding strategy that transforms grid cell activity from one (part of an) environment to another in a predictable manner (e.g. by aligning to a geometric axis in the environment), a possibility supported by recent work^32,33^. Similarly, a related possible cause of a shared representational geometry is input from the head direction (HD) system, which is expected to be “similarly different” across animals for the initial parts of the left and right trials in our data. Specifically, whatever HD cells are active for “left” trials would explicitly not be active for “right” trials due to the ring attractor organization of the HD system^34^. Although the hippocampal place cells in our data set do not behave in this literal exclusive-or pattern expected for head direction cells (we do not see negative left-right cell-by-cell correlations in Figure 5b), similarities in head direction across identically oriented compartments result in overlapping place cell maps compared to differently oriented compartments^35,36^, suggesting that a common head direction cell input may contribute to the cross-subject relationship between left and right trials in our hippocampal data (if not masked by symmetric cells as in the Gupta data set). If this is indeed the case, then our cross-subject hypertransform method would be able to predict, say, the place map of compartment oriented along the 45°axis in Grieves et al. (2018)^36^ based on the 135°compartment and the relationship between the 45°-135° compartments in a different subject, whereas in the Spiers et al. (2013)^35^ experiment, where the compartments all share the same orientation, it would be difficult to outperform the identity prediction.

In general, hippocampal representational similarity is shaped by many factors including not only the geometry of the environment as mentioned above, but also the identity of specific cues and their spatial configuration, the passage of time, and the animal’s internal understanding of the task structure^10,37^. In the two data sets we examined here, left and right trials are associated not only with different locations in space, but also with different head directions (at least for the part immediately after the choice point), different room cues, and different textures and rewards (for the Carey data). Based on these data we do not yet know which of these factors are subject-specific versus generalizable across subjects -- in other words, to what extent these factors contribute to cross-subject prediction. Future studies that systematically manipulate these factors while holding the others constant will reveal under what conditions cross-subject predictions, including those for experiences the subject has not yet had, can be obtained. More generally, uncovering the ways in which hippocampal activity is non-random can ultimately inform how processes such as generalization and structure learning are realized in neural circuits.

## Acknowledgements

We thank Jim Haxby and Feilong Ma for technical advice on the hyperalignment procedure; Youki Tanaka, Caleb Kemere, and the participants of the 2018 Methods in Neuroscience at Dartmouth summer school for discussion; Neil Burgess, Nancy Kanwisher, and Sam Mckenzie for valuable comments and suggestions, and A. David Redish for the use of one of the data sets in this study. This work was supported by HFSP Young Investigator Award RGY0088/2014, NWO VENI grant 863.10.013 and NSF CAREER award IOS-1844935.

## Author contributions

HTC wrote computer code and performed data analysis, in part using tools developed by JRM and MvdM. HTC and MvdM jointly conceived the analyses and discussed the results, with suggestions from JRM. HTC and MvdM wrote the paper with comments from JRM.

## Declaration of Interests

The authors declare no competing interests.

## STAR Methods

### RESOURCE AVAILABILITY

#### Lead contact

Further information and requests for resources and reagents should be directed to and will be fulfilled by the lead contact, Matthijs A. A. van der Meer.

#### Materials availability

This study did not generate new unique reagents.

#### Data and Code Availability

The code used in this study is available from Zenodo (https://doi.org/10.5281/zenodo.5076221) and GitHub (https://github.com/vandermeerlab/hc_hyperalign/tree/v1.0.0). The “Carey” data set used in this study is publicly available from DataLad (http://datasets.datalad.org/?dir=/workshops/mind-2017/MotivationalT) and the Gupta data is available from A. David Redish on request.

### EXPERIMENTAL MODEL AND SUBJECT DETAILS

Four male Long–Evans rats, aged 4–8 months at the start of behavioral training, were used in the first of two data sets used in this study (“Carey” data), and four male Fisher-Brown-Norway hybrid rats, aged 7–10 months at the start of recording sessions were used in the second data set (“Gupta” data). Full subject details are described in van der Meer et al. (2017)^17^ and Carey et al. (2019)^18^ Gupta et al. (2010)^19^ and Gupta et al. (2012)^20^ for the Gupta data.

### METHOD DETAILS

#### Behavioral task

We used two different data sets containing ensemble recordings of hippocampal CA1 neurons in rats performing T-maze tasks.

In the Carey data set, rats performed daily sessions on a T-maze where they had free choice between left and right arms. Rats were alternately food- and water-restricted across days; the left arm resulted in food reward (five 45 mg pellets), the right arm resulted in water reward (~0.2 ml sucrose solution). Rats ran 15-20 discrete trials per recording session, with no less than 5 trials for the least preferred choice (left or right).

In the Gupta data set, rats performed daily sessions on a continuous Multiple-T maze with free choice between left and right arms. Food pellet reward (four 45 mg pellets) was available either by choosing left only, right only, or alternating between left and right; which reward schedule was in effect was determined pseudorandomly at the start of daily recording sessions. In addition, the reward schedule switched approximately halfway throughout the session.

#### Criteria for inclusion of data

Only sessions with at least 40 simultaneously recorded neurons were included, leaving 19 of 24 total sessions for analysis (range: 50-178 neurons per session) of the Carey data set and 14 of 42 total sessions for analysis (range: 41-101 neurons per session) of the Gupta data set.

*Both data sets* consist of left (L) and right (R) trials from the choice point to the first reward site; the analyses in this study are concerned with how the relationship between L and R trials is encoded in hippocampal ensemble activity. To avoid the possibility that neural activity on a common trajectory shared between L and R is the main predictor of L and R relationship, data from the central stem of the maze was excluded.

##### Data preprocessing

###### Preparation of input data

Both data sets were preprocessed to obtain two types of neural activity matrices that form the starting point for all analyses (Figure S2). The first and main data type is the “Q-matrix”, which describes binned firing rate over time for simultaneously recorded neurons *[nNeurons x nTimeBins]* and is used in all main analyses. The second data type is the “TC-matrix” (place turning curves) matrices of dimension *[nNeurons x nSpaceBins]* for Figure 4 and S5).

Since both data sets contain different numbers of L and R trials within a session, trials were first subsampled so that an equal number of L and R trials were used. Next, because trials differed in length because of variations in running speed, all trials were truncated to the last 2.4 seconds (the median time between passing the choice point and reaching the reward site) so that only after-choice-point data was used.

To obtain Q-matrices for L and R trials, binned firing rate matrices (time bin width: 50 ms) were created for individual trials, smoothed with a window size = 1 s, σ = 50 ms Gaussian kernel, and then averaged across trials within each session.

To obtain TC-matrices for L and R trials, spike firing data with only running speed > 5 cm/s data were averaged for each place bin (~3 cm per bin) across each session, then smoothed with a window size =11 bins, σ = 1 bin Gaussian kernel. Only data from the last 41 place bins were included so that only after-choice-point data was used.

###### Criterion for exclusion of interneurons (analyses used in Figure 5 and Figure S3)

Neurons with mean firing rate > 10 Hz across the entire recording session were classified as putative interneurons. These were excluded for the correlation analysis in Figure 5, because otherwise variations in firing rates between putative interneurons and projection neurons would dominate the population vector correlations (described below). In Figure S3a we verify that inclusion of interneurons was not required for the main results, and noted that source sessions whose cross-subject predictions were consistently worse than the others (for instance, row 2 of R1 in Figure 3a, top row) were sessions without putative interneurons.

###### Normalization (analyses used in Figure S3)

Normalization of the input data was conducted by dividing the L2-norm of each row (neuron) or z-scoring each row of data matrices, *independently* for the L and R parts of the input data matrices. Although it may seem intuitive to normalize each entire row of the input data (i.e. L and R data together), this actually introduces an artificial anticorrelation between the L and R parts of the matrix, such that even on data where no relationship exists between L and R, a relationship is introduced by normalization. Thus, we avoided normalization across L and R when testing cross-subject prediction. Note that normalization was only performed on data with putative interneurons removed (Figure S3).

##### Hypertransform analysis procedure

###### Overview

The overall objective of the hypertransform procedure is to predict R data in a “target” subject based on (1) the target subject L data, and (2) the L and R data of a different “source” subject (see Figure 1 for a complete description and schematic). Each step of the procedure is described in detail below.

Each recording session was used as source and paired with all sessions from all other subjects to form cross-subject source-target pairs (260 unique source-target session pairs for the Carey data and 142 pairs for the Gupta data).

###### PCA

After preprocessing the data as described above, principal component analysis (PCA; *svd* function in MATLAB R2018b) was applied to concatenated L and R neural data matrices to reduce to the same dimensionality because (1) there were unequal numbers of neurons recorded across sessions, and (2) to aid in generalization across sessions. Ten principal components (PCs), accounting for approximately 95% of the variance (Figure S6d) were then used to project L and R matrices into neural activity trajectories in each subject’s own PCA space.

###### Hyperalignment

Hyperalignment is a procedure that maps neural activity from a number (>1) of input sessions into a common space. It does so by iteratively finding the set of Procrustes transformations (one for each input session) that minimize the Euclidean distance between input trajectories in the common space based on rotation, reflection and translation (and scaling, in some implementations; we did not use scaling in this study). This procedure is commonly used to align fMRI activity trajectories in subject-specific spaces into a common representational space (Haxby et al. 2011)^11^. In our specific version of this procedure, we aligned L and R neural trajectories in the PCA spaces from a source subject and a target subject into a common space by using the *hyperalign* function in *hypertools^16^ (matlab version)*. This results in two matrices, one for each subject, that map each subject’s PCA space into the common space.

###### Hypertransform

In this common space, another Procrustes transformation was derived to find the mapping between the L and R neural trajectories of the source subject (as above, scaling was disabled). This transformation (hypertransform, HT) was then applied to the target subject’s L trajectory to yield a *predicted* R trajectory. Note that this transformation is not only subject-specific but *source-target-pair-specific* since the common space is unique for each source-target pair used.

###### Projection back into neural space

Next, the inverse of the subject-specific hyperalignment matrix (projecting from PCA to common space) was applied to project the predicted R trajectory back to the target subject’s PCA space. Principal components obtained earlier were used to reconstruct the predicted R trajectory in the PCA space into the predicted R data matrix (Q or TC) for the target subject.

###### Shuffles and associated metrics

The prediction error for a specific source-target pair is calculated by summing squared errors between predicted and actual R data matrices of the target subject and dividing by the number of neurons. This error is compared against a chance distribution by permuting (shuffling) the rows of the R data matrix of the source subject (but keeping the L matrix intact; **row shuffle**, see Figure S2b) and repeating the hypertransform procedure. This shuffle breaks any relationship between L and R activity at the cell-by-cell level. For example, if the relationship between L and R firing activity is random, then permuting the source R data matrix in this manner will make no difference in the ability to predict the target R data matrix. As a result, for random data (such as our “independent” simulation in top row of Figure 6a), shuffled predicted errors should not be different from the actual observed predicted error. Note that row shuffles can create idiosyncratic principal components not present in the original data, affecting the extent to which source and target sessions can be aligned. Because hyperalignment (when used on two sessions, as in this study) “anchors” to the source session, asymmetries in the z-scored error relative to shuffle can arise, i.e. source-target and target-source can have numerically different z-scores.

Next, the **shift shuffle** preserves the mean firing rates of L and R for each neuron, but breaks the temporal relationship between L and R by circularly shifting each row of the source R matrix independently by a length randomly chosen from [1, number of total bins] (while keeping the L matrix intact; Figure S2b). If cross-subject prediction based on the true data outperforms this shuffle, then the specific firing fields matter, above and beyond mean firing rate relationships between L and R.

For each shuffling method, 1000 shuffles were conducted for each source-target pair, resulting in 1000 shuffled prediction errors which were compared to the prediction error observed for the true data using three metrics: (1) z-scoring the actually observed error relative to the distribution of shuffle predictions (lower z-scores indicate better prediction), (2) raw error compared to the mean error of the shuffle distribution (lower observed error indicates better prediction), and (3) proportion of the shuffle distribution whose error was smaller than actual observed error (lower proportion indicate better prediction).

###### Identity transform (used in Figure 5)

To test if a better-than-chance cross-subject prediction is simply due to the similarity of L and R neural activity within the target subject, a within-subject prediction is obtained by using a duplicate of target subject’s L trajectory in the common space as the predicted R trajectory, i.e. making the L-R transformation equal to the identity transformation. This predicted trajectory was then used to reconstruct the predicted target subject’s R data matrix as in the hypertransform procedure.

###### Split-half prediction (used in Figure 3 and Figure 4 insets)

To compare the accuracy of cross-subject hypertransform prediction to the best possible prediction given the variability in the target R data, we obtained a split-half prediction by averaging across a randomly-chosen half of right trials from the target session, resulting in a “lower bound” prediction error.

###### Variation: withheld data (used in Figure 4)

To determine the importance of including the data to be predicted (target R) in the hyperalignment step, we performed a number of analysis variations withholding different amounts of R data. In the first variation (Figure 4a), we withhold all target R data. Specifically, only source and target L were hyperaligned into the common space, in which the source R data was projected from the PCA space using the source matrix obtained from L alignment between the two subjects. The source L-R transformation and the predicted R matrix were then obtained as in the default hypertransform procedure. (Replacing the target R data matrix with zeros before hyperalignment yielded similar results to complete withholding, so we report only complete withholding results.).

In the second variation, a randomly-chosen single right trial from the target session was used for hyperalignment, and the average of the remaining trials were withheld as target R to be predicted (Figure 4b). The rest of the procedure follows the above hypertransform procedure. The split-half prediction in this procedure was obtained from half of the withheld trials.

###### Variation: PCA only (used in Figure 3 and Figure 4)

To test whether a relationship between L and R already exists in subject-specific PCA spaces, we modified the hypertransform procedure as follows: instead of aligning neural activity trajectories in the common space through hyperalignment, a L to R linear (Procrustes) transformation was derived from the source subject’s PCA space and directly applied to the L trajectory in target subject’s PCA space to obtain the predicted R trajectory. This predicted trajectory was then used to reconstruct the predicted target subject’s R data matrix as in hypertransform procedure.

#### Simulations

As a first step towards understanding the possible explanation for cross-subject prediction, simulated neural activity matrices were generated using 1-D Gaussians with three parameters: time (the time bin where a neuron has its maximal firing activity; randomly chosen from [1, 48] out of 48 time bins), peak firing rate (FR; randomly chosen from [10, 20] Hz) and width (randomly chosen from [2, 7] bins standard deviation). Several scenarios were created to test the possibility that different potential place cell properties may account for the observed cross-subject prediction:

- **ind-ind**: Neurons have a fixed, independent probability (0.5) of having a 1-D Gaussian place field on L and/or R, with the three parameters of time, peak firing rate (FR) and width randomly and independently chosen for L and R.
- **x-or**: Neurons only have a field on *either* L or R but not both, and parameters of the field are chosen randomly as ind-ind
- **ind-same-all**: Neurons have a fixed independent probability of having L and R fields as in **ind-ind** but with the additional constraint that neurons with both L and R fields must have the same three parameters.
- **sim. HT**: The activity on L is simulated by assigning each neuron an independent probability of having a field whose parameters are randomly chosen, then the activity on R is obtained by applying L-R transform (hypertransform, HT) from real data (Carey) to the simulated L activity.
- **ind-same-time**: Similar to **ind-same-all**, neurons have a fixed independent probability of having L and R fields as in **ind-ind**, but if when a cell has fields in both, one of the three parameters: **time** is the same for L and R, and FR and width are independently chosen.
- **ind-same-FR**: Similar to **ind-same-all**, neurons have a fixed independent probability of having L and R fields as in **ind-ind**, but if when a cell has fields in both, one of the three parameters: **FR** is the same for L and R, and time and width are independently chosen.
- **ind-same-width**: Similar to **ind-same-all**, neurons have a fixed independent probability of having L and R fields as in **ind-ind**, but if when a cell has fields in both, one of the three parameters: **width** is the same for L and R, and time and FR are independently chosen.

For each simulation scenario, we generated a matching number of sessions (19) as in Carey data and a matching number of neurons within each session as a dataset. To avoid the possibility that one particular randomly generated dataset biases the results, 100 datasets were created and all statistics were averaged across datasets to match the real data.

For each of the 100 datasets in **sim. HT**, a randomly chosen session from the Carey data was used to hyperalign with simulated L activity (R activity was replaced with zeros first). The hypertransform obtained from the real session was then applied to the simulated L trajectory in the common space to obtain a simulated R trajectory. This simulated trajectory was then used to reconstruct the simulated R activity as in hypertransform procedure.

### QUANTIFICATION AND STATISTICAL ANALYSIS

All statistics were performed in MATLAB (The Mathworks, 2018b).

#### Correlation analysis of neural and simulated data

All correlation coefficients were computed using the MATLAB *corrcoef* function.

##### Cell-by-cell (row-wise) correlations

For each cell (a row in the neural data matrix), a correlation coefficient between L and R was computed (see schematic in Figure S2). Whitening noise (matrices of same size as L and R, in which each element is a number sampled from *Uniform*(0, 10^−5^)) was added so that a coefficient can be computed when there is no activity on either L and R trials, which would otherwise result in zero variance.

##### Population vector (PV; column-wise) correlations

L and R data matrices were (horizontally) concatenated, and a correlation coefficient was computed between each time point (a column) and all other columns of the L and R concatenated matrix (see schematic in Figure S2). Higher off-diagonal values (the values along the diagonal of the lower left quadrant) of this concatenated matrix indicate higher correlated ensemble neural activity between L and R.

For both cell-by-cell and PV correlations, results were combined across subjects by appending all cells and sessions into a pooled data set regardless of what subject they came from.

#### Comparison of the hypertransform prediction to control predictions

##### Comparison to shuffles

For the hypertransform prediction error (between predicted and actual neural activity) of each source-target pair, a number of measures were computed: a z-score against the shuffled distribution, the proportion of shuffles with lower error, and the raw error compared to the mean of the shuffle errors. Note that raw errors were normalized by dividing by the number of neurons in the target session so that the prediction accuracy does not depend on the number of neurons being predicted. These measures are shown for each individual source-target pair as an element in a matrix such as one in Figure 3 (top row). Then, results across all source-target pairs are combined and plotted as a histogram; all summary statistics (median +/− SEM across all unique source-target subject pairs) and significance tests (Wilcoxon signed-rank tests against zero) are performed on this pooled data.

##### Comparison to PCA-only (Figure 3 and S4) and identity (Figure 5) predictions

For each source-target pair, the hypertransform prediction error is compared to the ones obtained from PCA-only and identity predictions (described in *Identity transform* and *Variation: PCA only* sections respectively). Then, results across all source-target pairs are combined and plotted as a histogram; all summary statistics (percentage of unique source-target subject pairs whose HT predictions are better) and significance tests (binomial test) are performed on this pooled data.

#### Identification of place fields

For Figure S7 only, we identified neurons as having a place field if three consecutive bins had a firing rate >5 Hz,on either the left arm (L), the right arm (R), or both. Only neurons with fields on both arms (6.98% of all neurons) are included in the analysis investigating the systematic mapping between L and R in Carey data set and in hypertransform (HT) predictions (Figure S7).

## Supplementary Information

**Figure S1:**
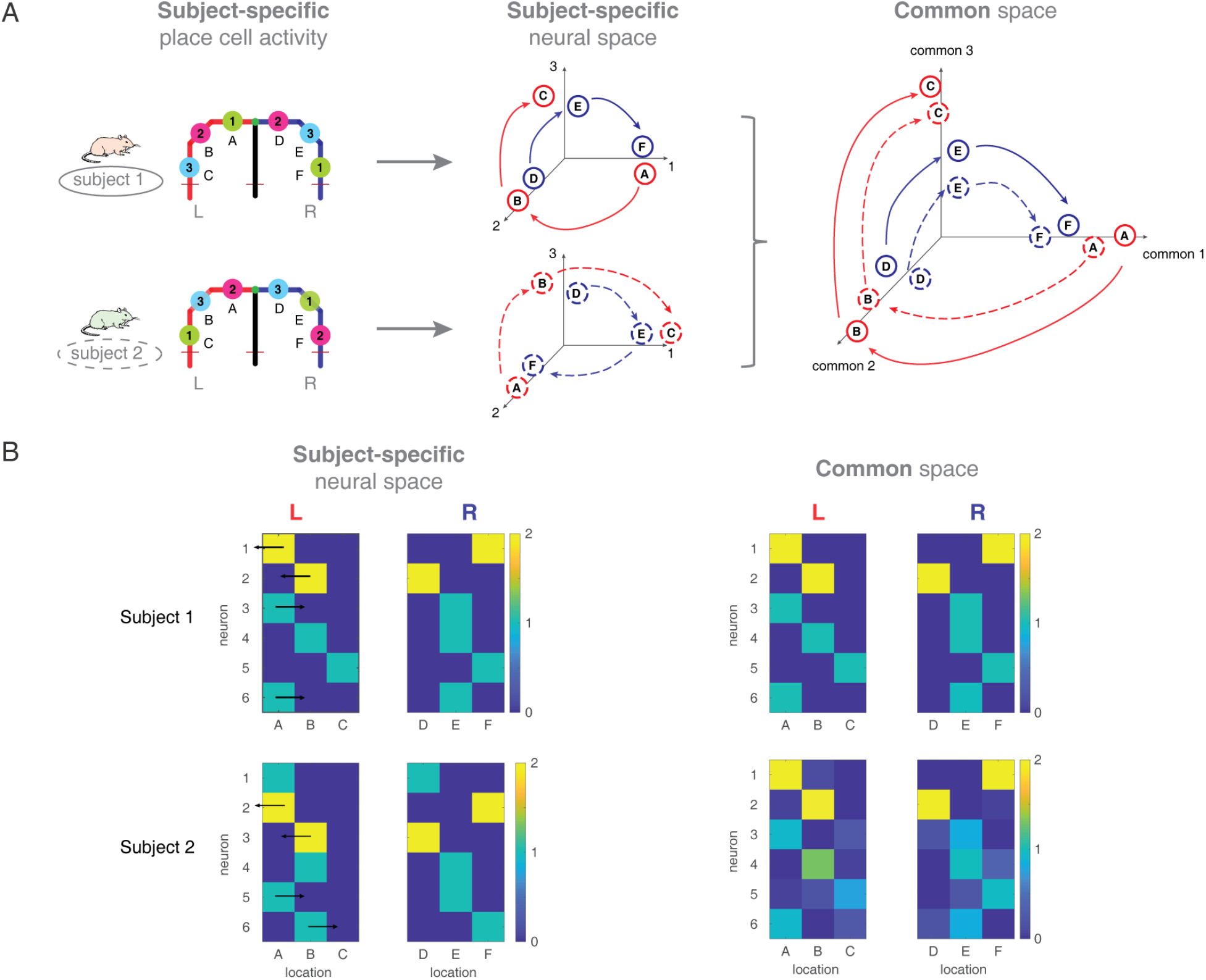
Schematic of hyperalignment applied to hippocampal place cell activity. **(A)** Schematic example of shared structure in place cells of two subjects on the left (L; locations A, B, C) and right arms (R; locations D, E, F) of a T-maze (left column). Individual neurons of different subjects (indexed by 1, 2, 3) have idiosyncratic place field locations for example, neuron 1 of subject 1 has place fields in A and F, whereas neuron 1 of subject 2 has fields in C and E (note that for visualization purposes, these neurons have place fields on both arms, but this is not a requirement for our analysis.) The ensemble neural activity during L and R runs can be plotted in a subject-specific neural space whose x-axis is the firing rate (FR) of neuron 1, y-axis is the FR of neuron 2, etc. (middle column; red line indicates trajectory of L and blue indicates R). Although subjects have different trajectories of L and R in their individual neural spaces, the trajectories for the two subjects can be made to line up by a rotation of the axes (e.g. rotate the x-axis of subject 1 to line up with subject 2’s y-axis). Hyperalignment finds the subject-specific transformation that minimizes the Euclidean distances of trajectories between subjects, and projects those neural activity trajectories into a common space (right column). In this common space, axes are shared between subjects, and trajectories of different subjects are now aligned to each other. Note that the common relationship between L and R in this example follows an unrealistically simple, fixed spatial mapping rule (the neuron firing at the start of L always fires at the end of R etc.), unlike what the real data suggests (Figure 5C, Figure S7). **(B)**: Minimal example of shared structure across subjects with essentially uncorrelated cell-by-cell and column-wise correlations that is more consistent with the data. The rule for simulating this example is as follows: neurons with high firing rates circularly shift their place fields to the left, whereas half of low-firing-rate neurons shift their fields to the right (arrows) and the other half stay fixed (left column, subject-specific neural space). Hyperalignment successfully captures this relationship by rotating, reflecting and translating subject 2’s data in neural activity space to minimize the distance to subject 1’s data. As a result of this step, neurons with similar co-activity patterns, and neurons whose activity changes in a similar way across subjects (regardless of where their firing fields might be) become aligned in common space: the two high-firing neurons align across subjects, as do the two low-firing neurons that don’t change, and the two low-firing neurons that do (right column, common space; note that subject 2’s subject-specific data can be recovered by inverting the transformation into common space). The shared structure in the actual data, which has many more neurons and more varied firing patterns, is more complex, and mapped in PCA space; this example, which does not use PCA, is intended to provide a visual illustration of how cross-subject activity patterns can exist in the absence of single-cell or population vector correlations.

**Figure S2:**
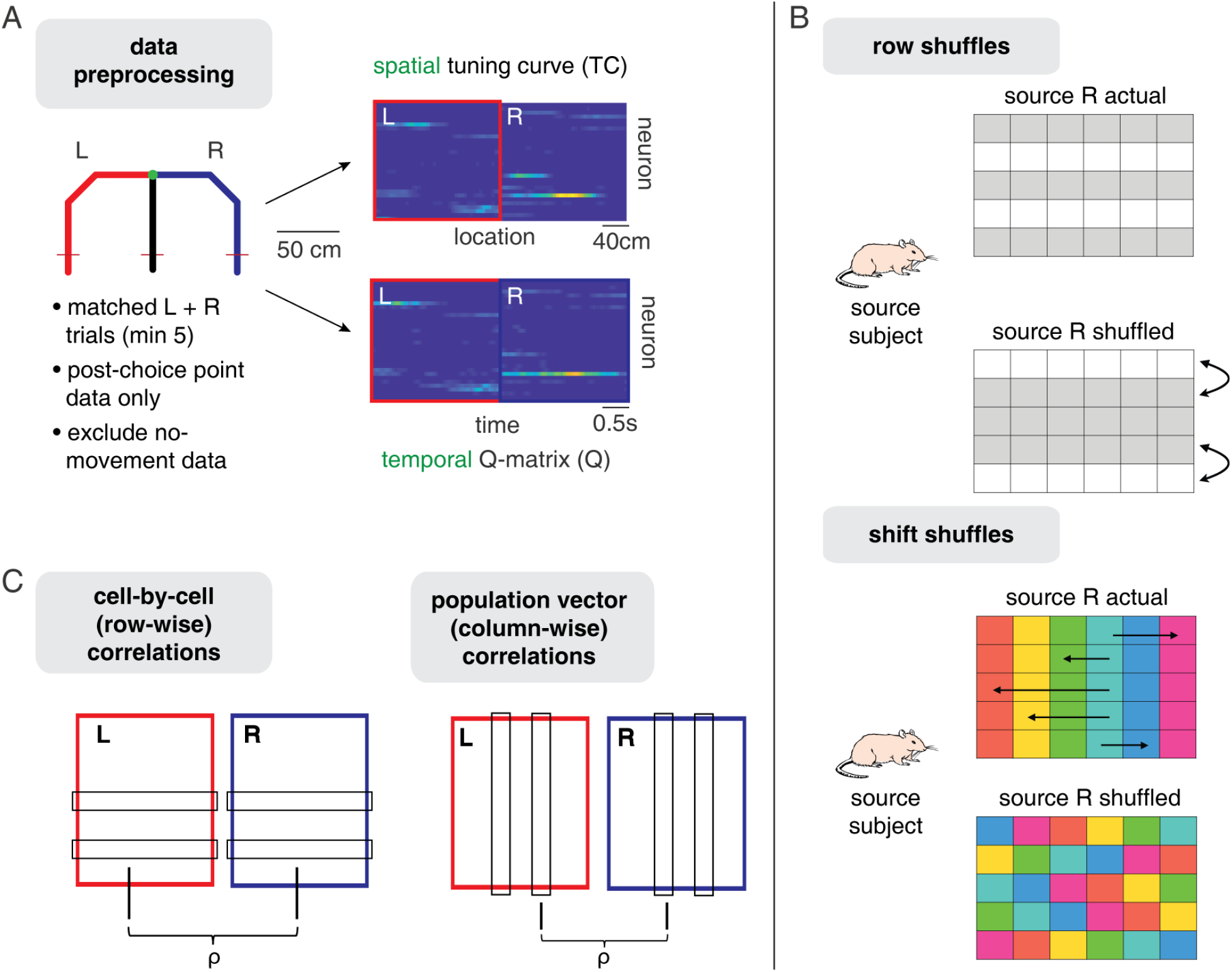
Data preprocessing and analysis schematics. **A:** Data preprocessing. Neural ensemble activity for left (L) and right trials (R) in each recording session was shaped into two types of input data matrices: the *Q-matrix*, which describes binned firing rate over time for simultaneously recorded neurons (dimension [nNeurons x nTimeBins]), used in the main analyses, and the *TC-matrix* (spatial tuning curves, dimension [nNeurons x nSpaceBins]). Trials were subsampled to obtain an equal number of L and R trials were used, and truncated to the last 2.4 seconds (the median time taken from the choice point to the end of a trial). Times when the animal was deemed to be stationary were excluded from analysis. **B**: Illustrations of the row shuffling procedure used in the main analysis (Figure 3) and shift shuffling procedure used in Figure 7. To obtain a distribution of chance cross-subject predictions, the analysis steps illustrated in Figure 1 were applied, except that for the “source” subject, the relationship between L and R activity was disrupted by either randomly permuting the rows of the R matrix (*row shuffles*) or circularly shifting each row of R matrix independently by a randomly chosen length (*shift shuffles*). **C**: Illustration of the correlation analyses used in Figures 5–6. *Cell-by-cell correlations* are obtained by row-wise correlating L and R activity for each cell, and then averaging across all cells. *Population vector correlations* are obtained by column-wise correlating activity at each time or location with activity at every other time or location. This yields one correlation matrix per session, which are then averaged across sessions.

**Figure S3:**
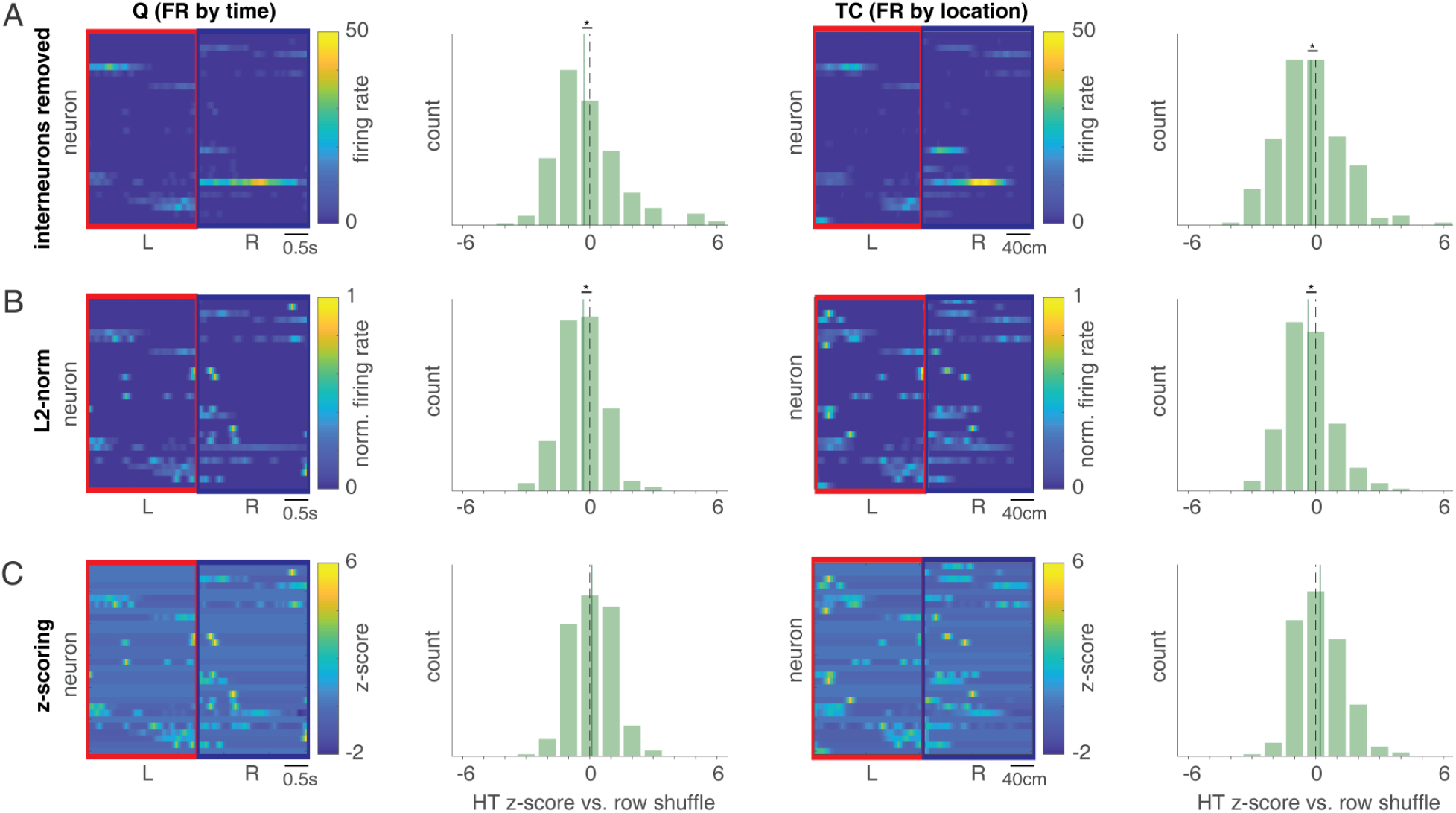
Cross-subject prediction is preserved when removing putative interneurons and normalizing L and R independently by L2-norm, but abolished when independently z-scoring L and R firing rates. **A**: Example L and R activity matrices with putative interneurons (mean firing rate > 10 Hz) removed, and corresponding cross-subject prediction z-score histograms compared to the distribution of shuffled predictions (z-scores lower than zero indicate better-than-chance prediction) for both temporal (**Q**) and spatial (**TC**) tuning curves. Both Q and TC show significantly better-than-chance cross-subject prediction (Q median: −0.42 +/− 0.46, SEM across unique source-target pairs, p < 0.01 for Wilcoxon signed rank test vs. 0; TC median: −0.34 +/− 0.43, p < 0.001 for HT vs. 0), indicating high-firing rate interneurons are not required for cross-subject prediction. **B**: Same layout as A, but L and R activity matrices were obtained by removing interneurons and dividing separately each L and R row by its L2-norm (vector length). Both Q and TC again show significantly better-than-chance prediction (Q median: −0.26 +/− 0.29, p < 0.001 for HT vs. 0; TC median: −0.43 +/− 0.32, p < 0.001 for HT vs. 0). **C**: Same layout as A, but L and R activity matrices were obtained by removing interneurons and z-scoring separately each L and R row. Unlike the L2-norm case, normalization by z-scoring does not show better-than-chance prediction; we speculate that this may be related to the relatively marginal performance of the L2-normed data compared to unnormalized data (compare panels **A** and **B**) combined with the tendency for the z-scoring especially to assign very large values to isolated pixels with a small but non-zero firing rate, introducing noise. Note that for both L2-norm and z-scoring cases, independently normalizing L and R activity (L separately from R), rather than normalizing L and R together (normalizing the entire row), avoids introducing artifactual anti-correlations between L and R which would be exploited by the cross-subject prediction algorithm even for independent data.

**Figure S4:**
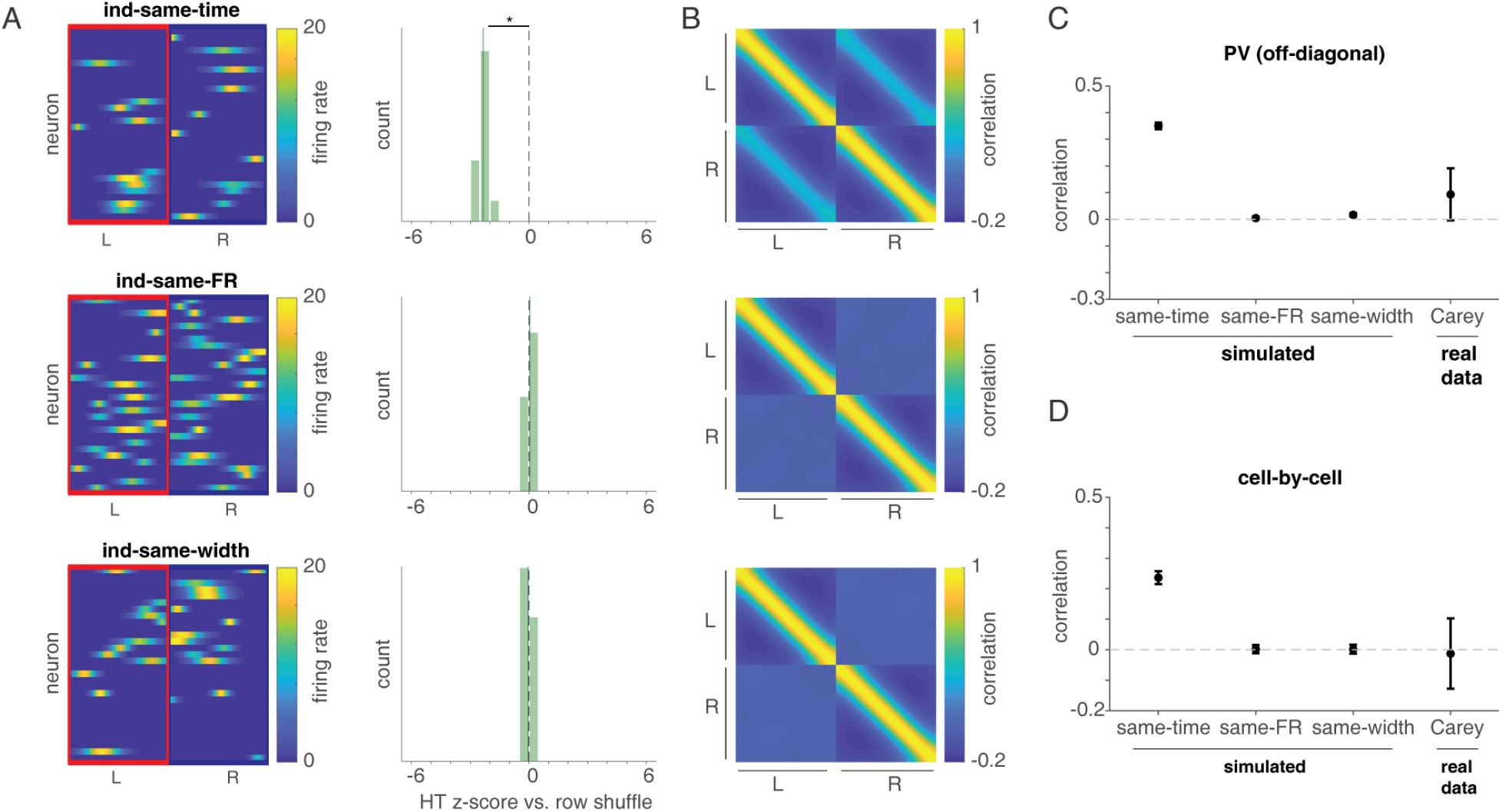
Simulations show that either the same place field firing time, peak firing rate, or place field width alone cannot account for cross-subject prediction in the Carey data. **A**: Example L and R activity matrices (left column) and histogram of z-scores of cross-subject prediction compared to the distribution of shuffle predictions (z-scores lower than zero indicate better-than-chance predictions; right column) of three simulated data sets. All three data sets are generated by assigning each neuron a fixed, independent probability (0.5) of having a field on L and/or R, but when a cell has fields in both, one of the three parameters of its 1-D Gaussian place field (**ind-same-time**, **ind-same-FR (firing rate)** or **ind-same-width**) is the same for L and R; the other two are randomly and independently chosen. **Ind-same-time** shows better-than-chance cross-subject predictions (median: −2.33 +/− 0.06, SEM across unique source-target pairs, p < 0.001 for Wilcoxon signed rank test vs. zero) but **ind-same-FR** and **ind-same-width** do not, indicating cross-subject predictions cannot be better than chance when time points of fields are unrelated between L and R. **B**: Population vector (PV; column-wise) correlations between ensemble activity at each time point and every other time point of the L and R activity matrices. **C**: Quantification of the median PV correlation between L and R (i.e. the values along the diagonal of the lower left quadrant in **B**). **Ind-same-time** shows highly positive correlation (median r = 0.35 +/− 0.01) since for every location of L where there are fields, there would some chance that some (although random) amount of ensemble activity would appear on the same location of R. In contrast, L and R in **ind-same-FR** and **ind-same-width** are nearly uncorrelated since no prediction of ensemble activity of one location on L can be made based on the same location of R (see also **ind-ind** in Figure 5). The **ind-same-time** case is potentially consistent with the Carey data as measured by PV correlations. **D**: Cell-by-cell (row-wise) correlations of L and R. Here, the **ind-same-time** correlations are higher than in the Carey data (median r = 0.23 +/− 0.02 vs. median r = −0.02 +/− 0.12 for Carey), indicating that this scenario does not accurately capture the source of cross-subject prediction in the data.

**Figure S5:**
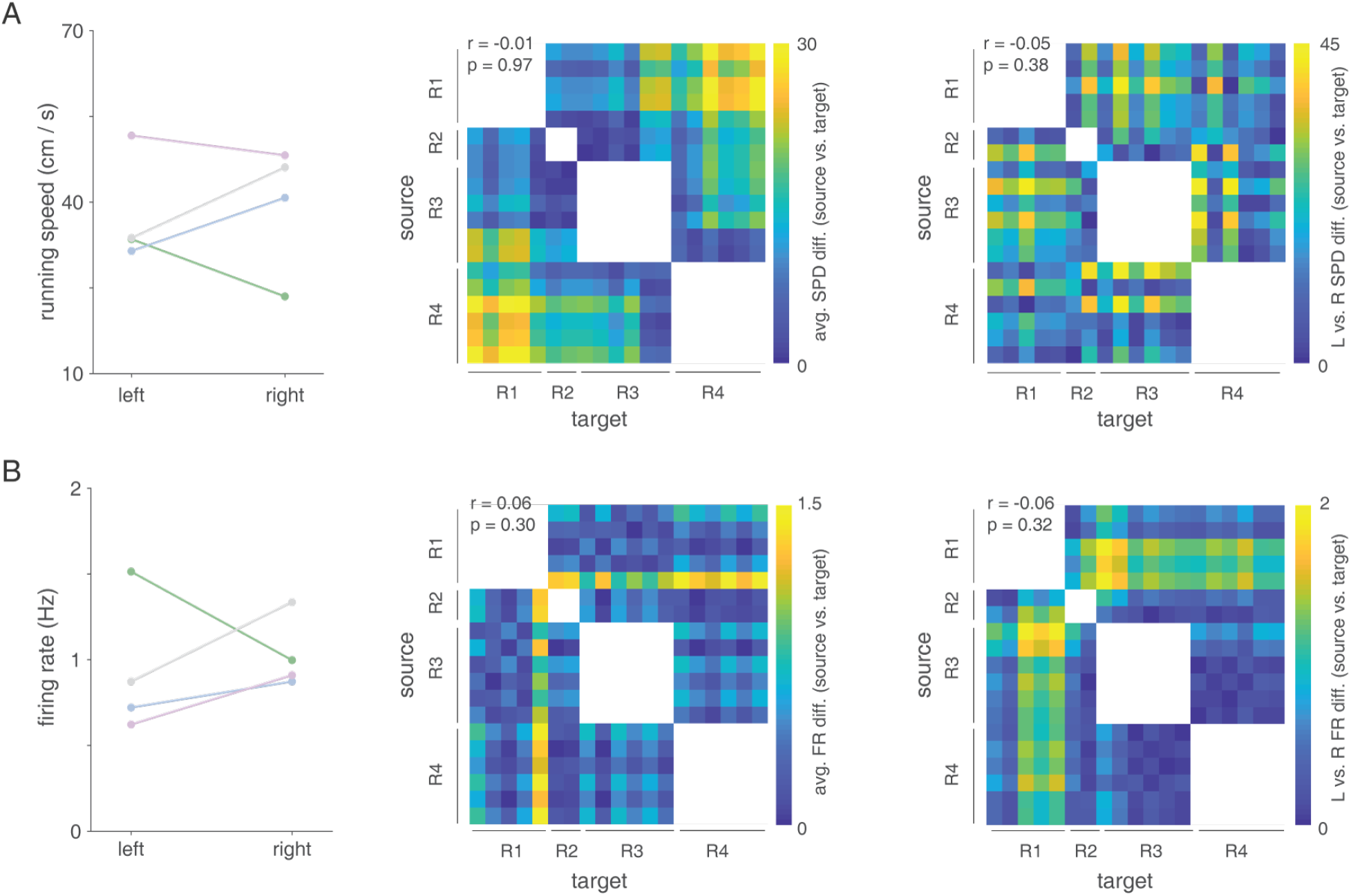
Differences in running speeds and firing rates between source-target subject pairs do not explain better-than-chance cross-subject hypertransform (HT) predictions. **A:** Average running speed across time points on both left trials (L) and right trials (R) for each subject (left column). There are significant differences between L and R (F_(1,14745)_ = 48.99, p < 0.001), between subjects (F(3, 14745) = 1406.95, p < 0.001) and the interaction of both (F_(3,14745)_ = 448.37, p < 0.001) in running speed. Given this variability in speed, we tested whether differences in average running speed between source and target sessions were related to prediction accuracy (HT z-score matrix in Figure 3a, top row), reasoning that more similar speeds may enable better predictions. Contrary to this expectation, we did not find a significant correlation between absolute running speed differences and prediction accuracy (r = −0.01, p = 0.97; the specific quantity computed was | speed_source_–speed_target_|). Since there is an interaction between L/R and subject, we also calculated spd-diff_session_ = |speedL_session_–speedR_session_| for each session and a specific quantity |spd-diff_source_–spd-diff_target_| for each source-target pair, capturing how similar (or different) L/R speed differences are between source-target pairs (right column). Again, we did not find L/R speed differences significantly correlate with the prediction accuracy (r = –0.05, p = 0.38). **B:** Average firing rates (FR) across time points and neurons on both left trials (L) and right trials (R) for each subject (left column). There are also significant differences between L and R (F_1,181624_) = 16.13, p < 0.001), between subjects (F_(3,181624)_ = 178.16, p < 0.001) and the interaction of both (F_(3,181624)_ = 132.31, p < 0.001). Again, we repeated the analysis in **A** and did not find significant correlations between the prediction accuracy and the absolute difference in average FR between source-target pairs (r = 0.06, p = 0.30), and between the prediction accuracy and the absolute L and R difference in FR between source-target pairs (r = −0.06, p = 0.32).

**Figure S6:**
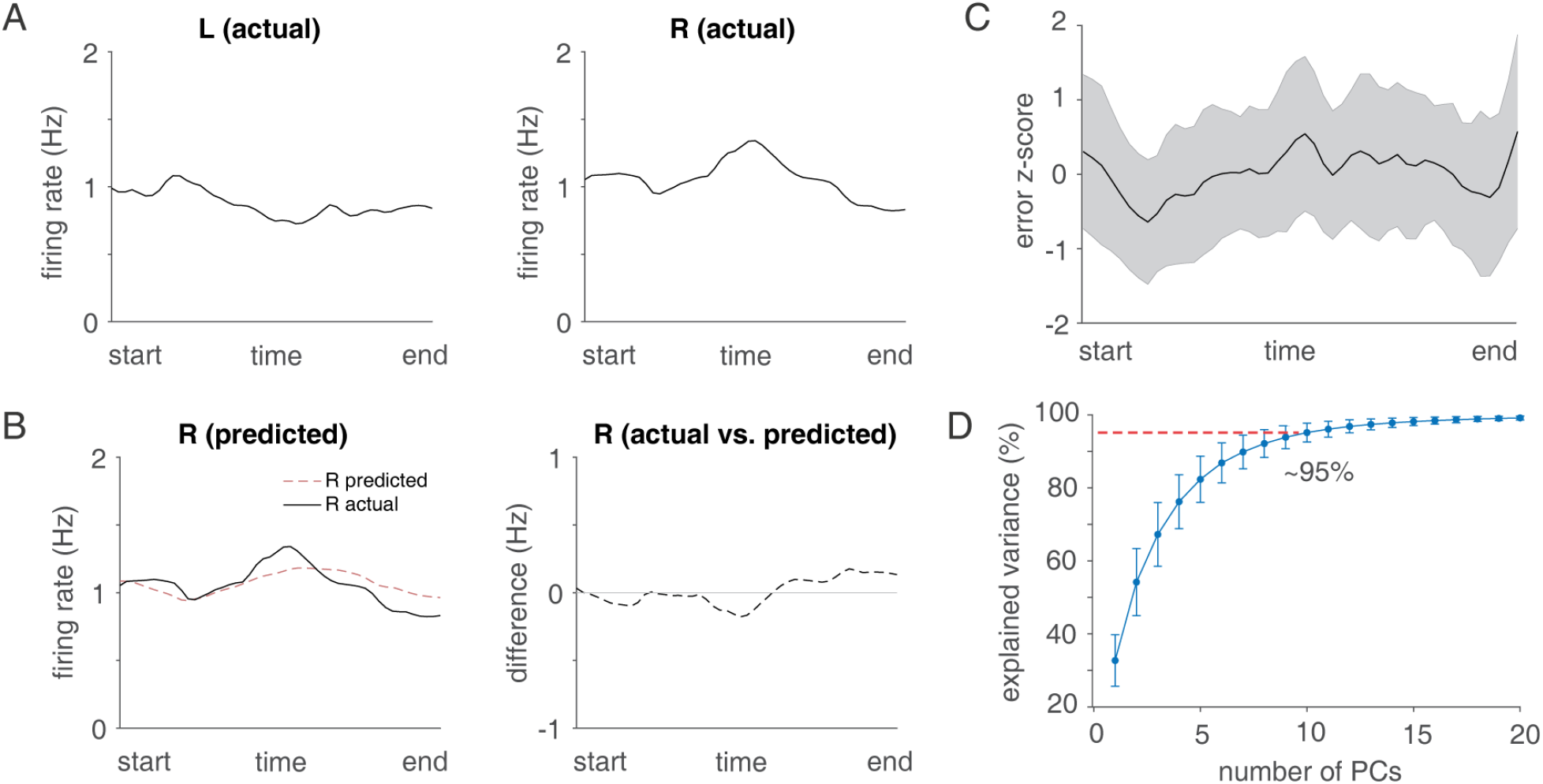
Cross-subject hypertransform (HT) prediction is not due to particular time points such as the start or end of the trial. **A:** Average firing rates (FR) across neurons for each time point for both left trials (L) and right trials (R) in the Carey dataset. **B:** Comparison of average firing rate across neurons over time with average predicted firing rate. Although FR of both L and R varied across time points, we did not find any time point at which FR is substantially high or low, which could potentially be exploited by the HT prediction, for example a high activity at the start of L is always mapped to the end of R. Similarly, the predicted R activity and its difference with actual R also show varying patterns but not a particular time point is highlighted. **C**: Z-scores of cross-subject prediction errors (normalized within each session) as a function of time, averaged across sessions. Errors varied across time but did not highlight particular time points, indicating that cross-subject prediction is not disproportionately due to certain time points such as the end of the trial. **D:** Explained variance as a function of the number of principal components (PCs). In our hypertransform procedure, 10 PCs were used, which accounts for ~95% of the variance of data.

**Figure S7:**
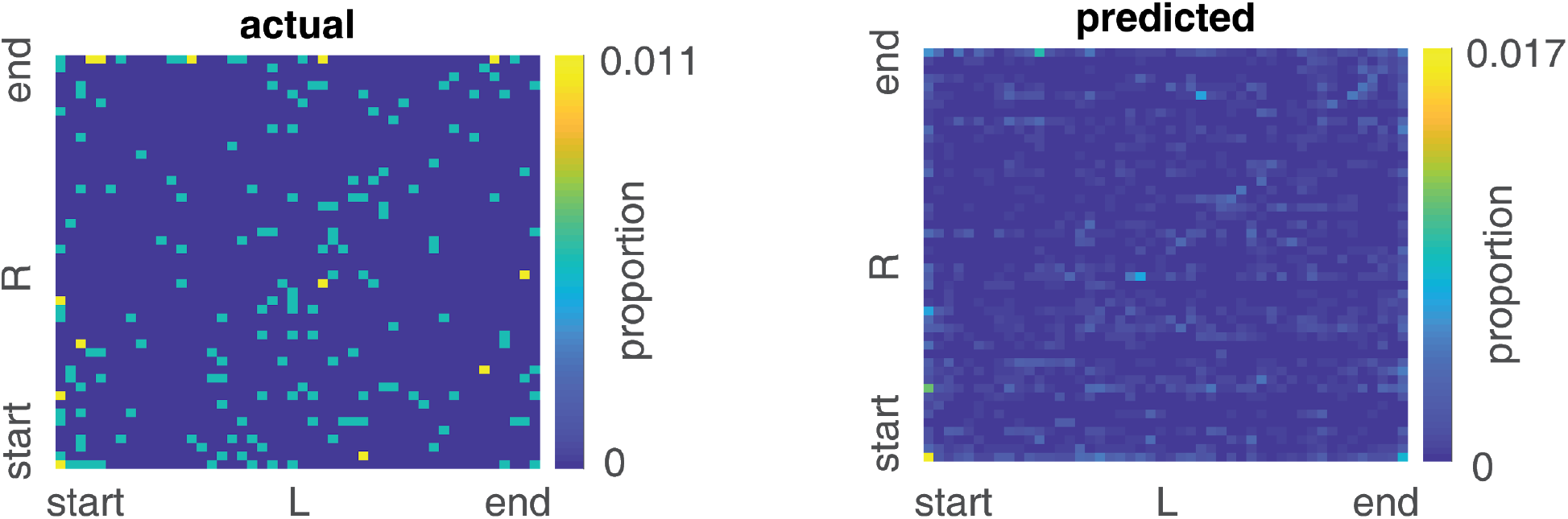
No obvious single-cell mapping between left (L) and right (R) trials for both actual and predicted data. To investigate if there is a systematic mapping between in L and R in actual data and hypertransform (HT) predictions, we plotted temporal field indices of L and R for neurons which have fields on both (6.98% of all neurons) on a matrix where x-axis are temporal field indices of L and y-axis are indices of R (see *Methods* for how fields were detected). If a neuron has a temporal field on the end of L and a field on the start of R, it is counted as 1 on the bottom right corner of the matrix, and total counts are divided by the number of neurons included. We did not observe a simple rule characterizing mapping in temporal fields between L and R in actual data; for example, it is not the case that neurons at the end of L are systematically being mapped to the start of R. Other than some subtle identity predictions (along the diagonal), we did not find an obvious single-cell mapping between L and predicted R, indicating that the HT prediction captures ensemble-level relationships rather than single cell relationships.

## Notes

### Competing Interest Statement

The authors have declared no competing interest.

### Summary of Updates

Minor pre-publication edits.

